# Adaptation to free-living drives loss of beneficial endosymbiosis through metabolic trade-offs

**DOI:** 10.64898/2026.04.16.718893

**Authors:** Erika M. Hansson, Irma Vitonytė, Guy Leonard, Rachel Smith, Daniel Malumphy Mon-tesdeoca, Fiona R. Savory, Perdita E. Barran, Luke T. Dunning, Jon Slate, Thomas A. Richards, Andrew P. Beckerman, Duncan D. Cameron, Michael A. Brockhurst

## Abstract

Symbioses are widespread (1) and underpin the function of diverse ecosystems (2–6), but their evolutionary stability is challenging to explain (7,8). Fitness trade-offs between contrasting intracellular and extracellular niches could act to stabilise endosymbioses because adaptation to either niche is predicted to reduce fitness in the alternate niche, thus reinforcing symbiosis (8,9). Here, we experimentally evolved four diverse *Chlorella* green algal endosymbionts of *Paramecium bursaria* to free-living conditions supplying either an amino acid, as provisioned by hosts (10,11), or nitrate, as available in freshwater (12), as the sole nitrogen source. Experimental algal populations adapted to free-living environments, generally increasing in population density and cellular chlorophyll content over time. In one of the four endosymbiont strains, adaptation to the nitrate free-living environment, but not the amino acid environments, drove the loss of fitness benefits to the host in reconstituted symbioses. This loss was not associated with reduced ability to grow on host-provisioned amino acids, nor lost ability to release the sugars provisioned to the host (10,13). Genome sequencing of evolved algal lines revealed genomic divergence between nitrate-adapted and amino acid-adapted lines, affecting genes involved with metabolic organisation and intracellular resource transport. Untargeted metabolomic profiling further showed extensive changes to membrane remodelling and turnover in N-evolved lines. Together, our data support a role for metabolic trade-offs driving divergence between contrasting intracellular and extracellular niches, with nitrogen as a key environmental axis driving divergence. Fitness trade-offs may, therefore, be a general, simple mechanism acting to reinforce symbiosis, contributing to evolutionary stability.

## Main text

*Chlorella* green algae form a facultative endosymbiosis with the ciliate *Paramecium bursaria*, wherein the ciliate hosts hundreds of algal symbionts that it provides with organic nitrogen (amino acids) (10,11) in return for the products of photosynthesis (maltose and oxygen) (14). The species of *Chlorella* that form this symbiosis can also be found free-living in freshwater environments where (15), in contrast to the intracellular niche, inorganic nitrate is the most abundant source of nitrogen (12). We hypothesized that this contrast in nitrogen provisioning between intracellular and extracellular niches may mediate metabolic trade-offs driving specialisation into symbiotic versus free-living types (9). To test this, we experimentally evolved four genetically and geographically diverse endosymbiotic *Chlorella* strains under free-living conditions supplying either arginine (A-evolved) or glutamine (G-evolved), predicted to be supplied to endosymbionts by hosts (10,11), or nitrate (N-evolved), as available in freshwater, as the sole nitrogen source. Replicate populations were propagated by weekly serial transfer to fresh media for 30 weeks (approximately 50—150 generations). Except strain Symb_21 in nitrate, all lines increased in population density over time, indicating environmental adaptation, albeit at different rates per nitrogen source for some strains (Fig 1A; Table S1). Similarly, the per cell green fluorescence generally increased (Fig 1B, Table S1) suggesting higher investment in chloroplast and chlorophyll production, again at different rates per nitrogen source and strain combination. Some lines also increased in cell size, while patterns for cell granularity varied strongly between strains (Fig S1, Table S1).

**Figure 1:**
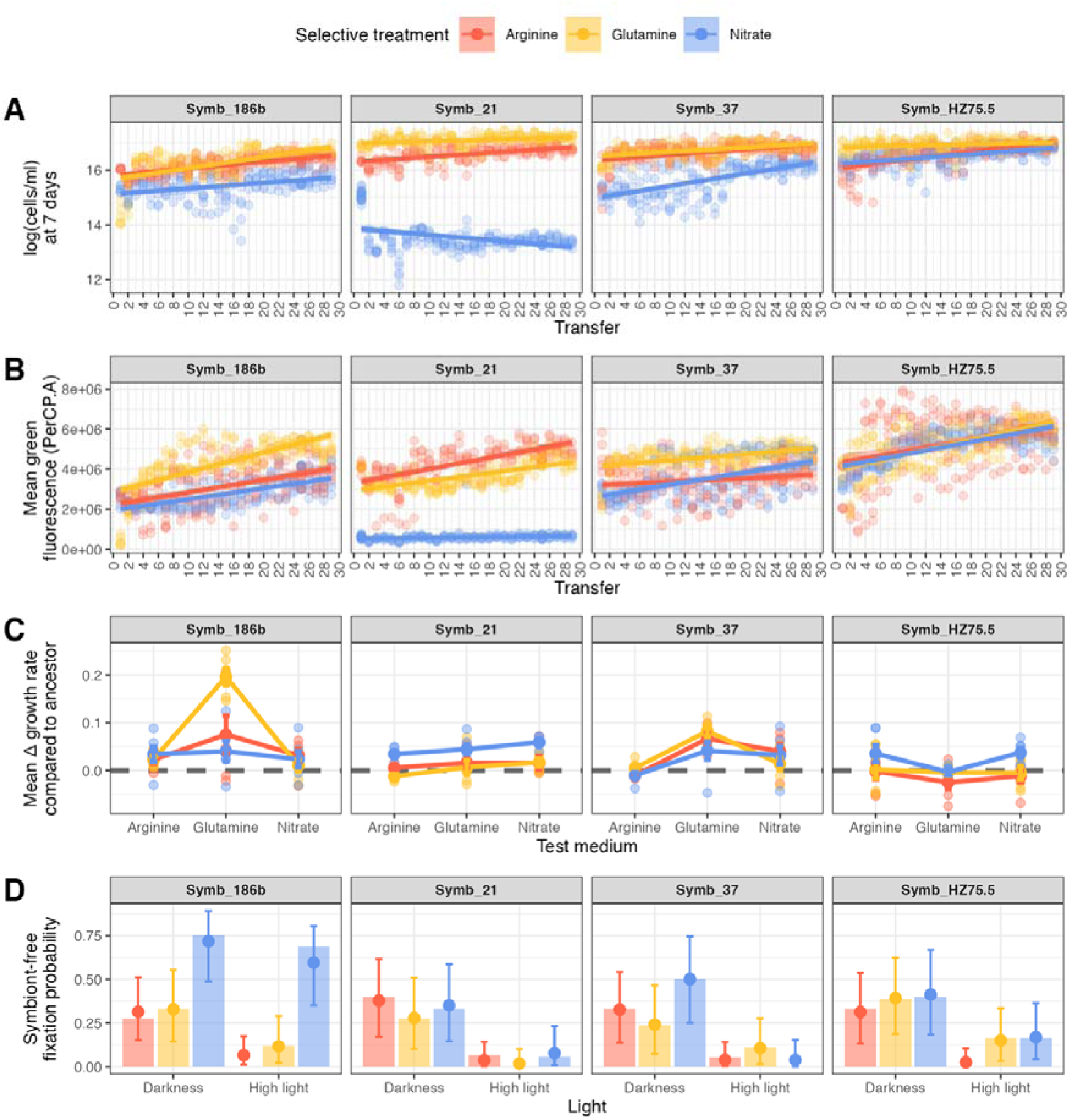
Phenotypes for the evolved algal strains in arginine (red), glutamine (yellow) and nitrate (blue). A) Day 7 population density of each strain across transfers. Raw data are shown as semi-transparent points, the solid line showing the linear trend with standard error ribbon. B) Day 7 mean green fluorescence of each strain across transfers. C) Change in growth rate of evolved strains compared to ancestor across different nitrogen sources. Raw data are shown as semi-transparent points with means and standard error represented as opaque points and error bars. D) Probability of the evolved strain losing to white cells in competition assays in darkness and high light. Proportion of populations where white cells became fixed is shown as bars. Posterior estimates with 95% CI from the Bayesian generalised linear model fit (Bernoulli likelihood, logit link) are shown as points and error bars.

To test adaptation to the nitrogen environment, growth rate of all lines in all three nitrogen sources was measured. We found that all strains grew as fast or faster in all nitrogen sources compared to their ancestors, with the largest increase not always being in the nitrogen source matching their selection background (Fig 1C; Table S2). This suggests an overall pattern of generalist adaptation to the free-living niche regardless of nitrogen source, with the largest increases in growth rate seen in A– and G-evolved Symb_186b and Symb_37 growing in glutamine (Fig 1C; Table S2).

Next, to test how free-living adaptation had impacted symbiosis, we used evolved algae from the endpoint of the serial transfer selection experiment to infect a common-garden *P. bursaria* host genotype, HA1 (16), which had previously been cured of its algal endosymbiont. We quantified the fitness effect for the host of carrying evolved algal endosymbionts by directly competing symbiotic HA1 against isogenic symbiont-free HA1 in darkness or light, where algal symbionts would be expected to be costly versus beneficial, respectively (16). Due to appreciable incidence of fixation of symbiont-free HA1 among replicates (e.g. Figure S2), we use the fixation probability of symbiont-free HA1 as a proxy for fitness, where a higher fixation probability indicates lower symbiotic fitness. As expected, light increased the fitness of symbiotic relative to non-symbiotic HA1, however the effect of prior nitrogen environment on symbiotic host fitness varied according to algal strain. Whereas prior nitrogen environment had no effect on symbiotic host fitness for two of the algal strains (Symb_21; Symb_37), A-evolved Symb_HZ75.5 saw a slightly increased symbiotic fitness for hosts, and for Symb_186b we observed strongly reduced symbiotic fitness for hosts carrying the N-evolved compared to the amino acid-evolved algae both in light and in darkness (Fig 1D; Table S3). As N-evolved Symb_186b strains showed increased growth in arginine as well as glutamine, it is unlikely that the loss of the symbiotic phenotype is directly explained by im-paired utilisation of host-provisioned amino acids. To test whether altered symbiotic ability could alternatively be explained by loss of algal sugar export, we quantified maltose and glucose export in the evolved Symb_186b lines in acidic pH outside the host. This mimics acidification of the perialgal vacuole which is a host signal to stimulate algal sugar release (17,18). N-evolved lines exported less maltose and glucose than A-evolved lines, but as much maltose and more glucose than the G-evolved lines, suggesting that loss of sugar export capacity did not explain loss of symbiosis (Fig S3; Table S4).

To understand the genomic basis of the observed phenotypic divergence of Symb_186b between the nitrate and amino acid treatments, we whole genome sequenced evolved Symb_186b lines from the endpoint of the selection experiment. Variants were called within genic regions against the ancestral reference genome (19). Overall, we observed 1516 fixed variants (i.e., with an allele frequency >= 95%), with no difference in the total number of variants per line between treatments (Fig S4; ANOVA: F = 0.1041; D.F. = 2, 15; p = 0.9). Most variants were within introns (98.4%; Fig S5) and are thus likely to modify gene regulation. There was strong genome-wide parallelism within treatments, with significant divergence between treatments at both the SNP (Δ distance = 0.14, 95% CI = [0.12, 0.14], *p* < 0.0001) and gene (Δ distance = 0.14, 95% CI = [0.13, 0.14], *p* < 0.0001) levels. To identify divergently evolving genes, we identified variants most strongly associated with each treatment, and retained those that were treatment-specific, mutated in at least two replicate populations per treatment and robustly called using a range of allele-frequency thresholds, resulting in a list comprising 216 variants of 168 unique genes (Supplementary materials 2). As such, these represented genes that evolved divergently between treatments and in parallel within treatments.

To gain an integrated overview of the functions affected by divergent parallel evolution we used gene ontology (GO) enrichment analysis, which revealed clear functional divergence between treatments (Fig 2). G-evolved lines were enriched for mutations affecting amino acid biosynthesis, ribosome assembly, and protein-RNA complex organisation (Table S5), indicative of elevated biosynthetic and translational activity. Parallel-evolving genes included those encoding glutamine synthetase, tRNA synthesases, ribosome-associated proteins, and AAA+ ATPases (Supplementary materials 2) involved in macromolecular assembly (20–22). In addition, G-lines were enriched for mutations affecting oxidative stress response and plastid-associated genes (Table S5), consistent with dynamic management of redox balance (e.g., peroxidases, glyoxalases (23,24)) and altered chloroplast structure and function (fibrillins (25)). A-evolved lines were enriched for mutations affecting transmembrane transport processes, particularly organic acid, dicarboxylate, and small molecule transport, alongside regulation of metabolic and biosynthetic processes (Table S6). Parallel-evolving genes included those encoding putative succinate/acetate transporters (e.g. SatP-like proteins), major facilitator superfamily transporters, and cation/proton exchangers (Supplementary materials 2), consistent with altered metabolite exchange across compartments (26–28). Other enriched functions included nitrogen assimilation and amino acid metabolism (Table S5), with parallel mutations in genes encoding carbamoyl-phosphate synthase and asparagine synthase, alongside transcriptional and chromatin regulators (Supplementary materials 2), indicating selection for altered metabolic flux (29–33). N-lines, by contrast, were enriched for mutations affecting protein catabolic processes, proteasome function, and organelle organisation (Table S5), suggesting altered protein turnover and cellular remodelling. Parallel-evolving genes included those encoding proteasome subunits, ubiquitin ligases, and a likely F-box superfamily gene, alongside lipid metabolic enzymes such as thiolases and alpha/beta hydrolases (Supplementary materials 2), further indicating altered protein and lipid recycling (34–38). Additionally, enriched functions included transport processes (e.g. ABC transporters and amino acid transporters; Supplementary materials 2) and stress-related functions (Table S5), further supporting a shift toward internal resource scavenging and cellular maintenance (39–41).

**Figure 2:**
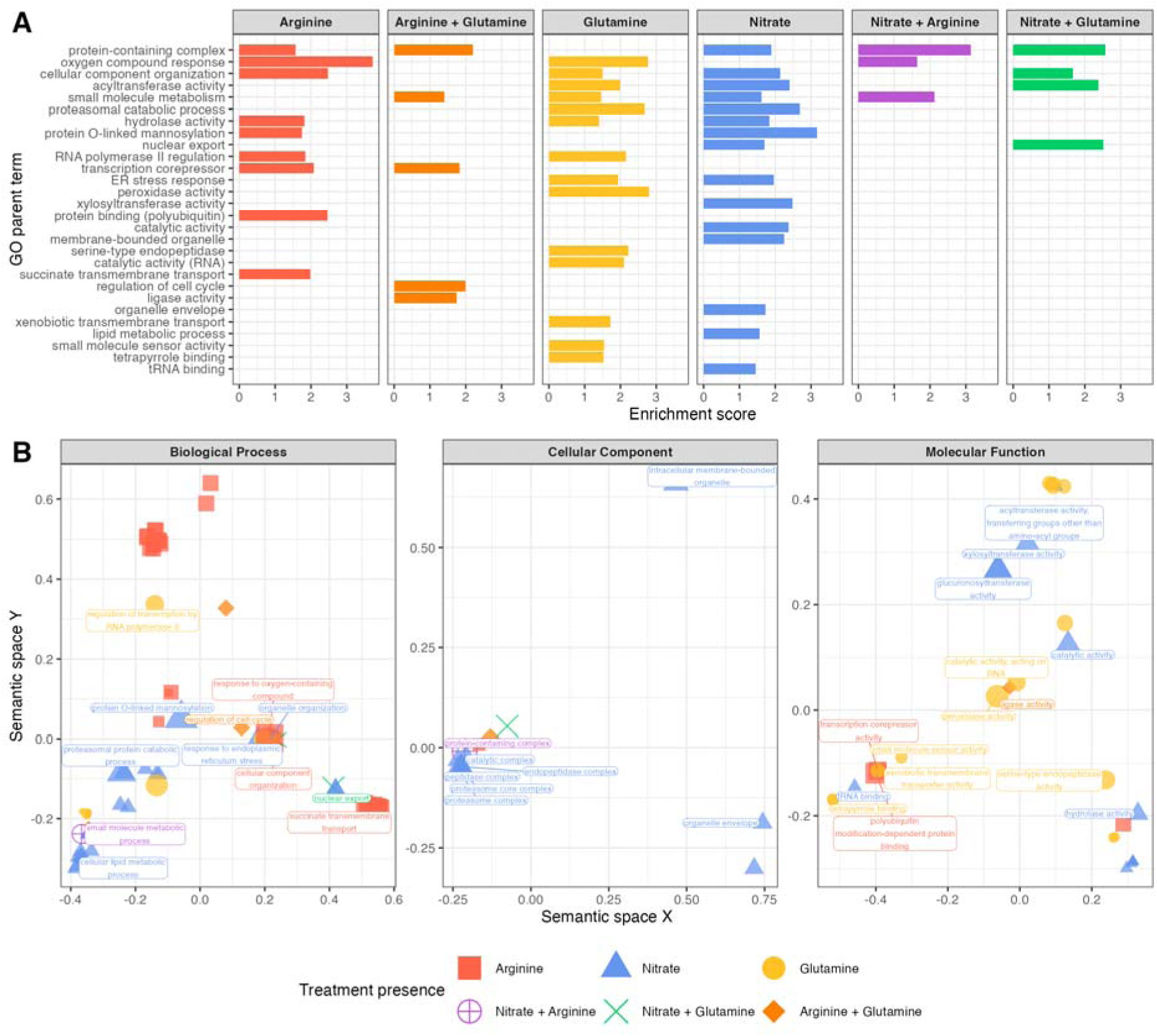
Top enriched GO terms in genes of interest shortlist by A) top parent terms and B) separated in semantic space. Bar length and point size is scaled by –log10(p-value) (i.e. larger bars/points = smaller p-value). Points are coloured by treatment presence and labelled with top associated GO terms, with priority in plot B given to labels associated with N-evolved lines.

Given the divergent genomic evolution observed in metabolic functions, and in particular the enrichment of lipid metabolism in the N-associated parallel mutations, we obtained untar-geted small molecule metabolomic profiles for the Symb_186b lines at five timepoints (transfers 1, 5, 10, 20, 30) using Desorption Electrospray Ionisation Mass Spectrometry (DESI-MSI), which is suited for detecting lipid composition changes. This revealed strong divergence of metabolomes between treatments (Fig 3, PERMANOVA F = 20.3; DF = 2, 84; p < 0.001). Top m/z peaks from PCA loadings and a random forest classifier were used to identify putative treatment-associated metabolites. Linear mixed models were used to determine whether treatment had a significant effect on the m/z peak intensity through time, resulting in a shortlist of 142 m/z peaks, of which 22 could be identified to at least class level with MS/MS (Fig S6, Table S6-8). A– and G-evolved lines were principally enriched for lipids involved in thylakoid membranes (42–44) and phosphoinositide-mediated signalling and membrane trafficking (45,46) or putatively mitochondrial membranes (47), respectively. By contrast, whereas N-evolved lines also showed elevated levels of thylakoid membrane-associated and signalling lipids, they were uniquely enriched for lipids involved in membrane remodelling and turnover (α-linolenic acid and lysophasphatidylglycerols (43,48)).

**Figure 3:**
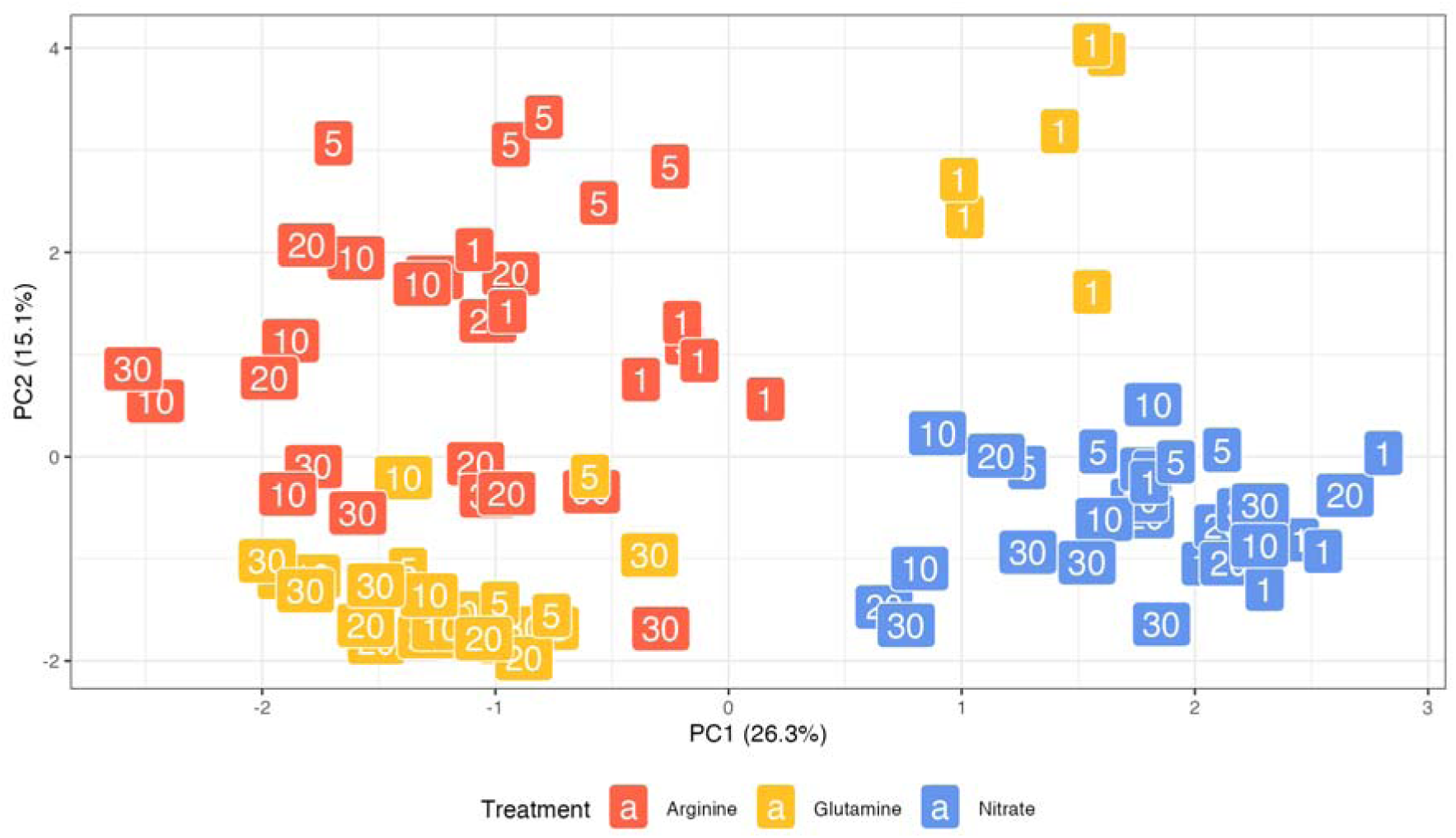
PCA for negative mode m/z peak intensities. Numeric labels show sampling week (1—30) during the selection experiment.

Fitness trade-offs between intracellular and extracellular environments have been proposed as a general stabilising mechanism for endosymbiosis because adaptation to each niche leads to loss of fitness in the other (8,9). Here, we provide direct experimental evidence for metabolic trade-offs centred on nitrogen utilization driving such divergence in the common freshwater microbial symbiosis between *P. bursaria* and *M. conductrix*. Algal endosymbiont adaptation to free-living within environments supplied with inorganic nitrogen (nitrate), but not those supplied with organic nitrogen (amino acids), drove the breakdown of symbiosis with hosts. N-evolved lines gained parallel mutations across a distinct range of functions versus A– or G-evolved lines, including protein degradation, lipid metabolism and internal nutrient scavenging. Moreover, N-evolved lines restructured their lipid profiles, suggesting divergence of membrane turnover and lipid signalling functions. Together, these patterns suggest the constraints of assimilating a particular form of nitrogen drove distinct metabolic strategies. Glutamine, a reduced nitrogen source, can be readily incorporated into central metabolism, providing both nitrogen and carbon skeletons with minimal energetic cost (49,50). Arginine, while also reduced, requires more complex, compartmentalised metabolism and redistribution of carbon (51,52). By contrast, nitrate must be reduced to ammonium prior to assimilation, a process requiring both substantial photosynthetically derived reductant and carbon skeletons for downstream amino acid biosynthesis (49,53). Accordingly, given this more stringent metabolic niche, N-evolved lines appear to have extensively remodelled their metabolism for efficient internal recycling of cellular resources, notably of proteins and lipids. Given that N-evolved lines did not entirely lose metabolism of host-provisioned amino acids, these data suggest that the loss of beneficial symbiosis we observe is likely caused by a mismatch between the N-evolved metabolic state and the within-host intracellular environment. Within the host, lower irradiance and organic nitrogen supply are likely to disrupt the balance selected for in the N-environment, simultaneously altering the carbon-nitrogen coupling and removing a major electron sink. Under these conditions, continued high rates of photosynthetic electron transport could lead to over-reduction and increased reactive oxygen species, while elevated protein and membrane turnover may impose additional energetic costs and destabilise the sustained metabolite exchange required for stable, beneficial endosymbiosis.

Previous studies show that newly established endosymbionts adapt to the intracellular environment through regulatory rewiring and metabolic reorganisation (15,54–57). Here we show that the converse also holds: Endosymbionts can rapidly lose symbiotic competence as they adapt to the free-living extracellular environment. The strong genetic parallelism observed across replicate evolving lines indicates that these changes are driven by selection rather than drift and cause consistent shifts in metabolic strategy dependent upon the nitrogen source. These results highlight the importance of selection-driven metabolic trade-offs between intracellular and extracellular niches, suggesting that adaptation to one environment can drive loss of performance in the other. Together, such opposing selective pressures are likely to reinforce symbiosis by constraining the existence of metabolic states compatible with both lifestyles.

## Methods

### Isolation of symbiotic algal strains

We isolated four algal symbionts from lab strains of host *Paramecium bursaria*: two *Chlorella variabilis* strains (Symb-1660/21-IV from host 1660/21, and Symb-HZ75.5-VI from host HZ75.5) and two *Micractinium conductrix* strains (Symb-1660/37-III from host 1660/37, and Symb-186b-X from host 186b). Prior to isolation, *P. bursaria* cultures were maintained in standard *Paramecium* growth conditions at 20°C under 12 µE m^-2^s^-1^ light (10:14 L:D cycle) in modified NCL medium (58) with 0.25 g Protozoan pellet (CBA053, Blades biological LTD) replacing the cereal grass leaves.

Symbiotic algae were isolated by washing *Paramecium* cells with NCL with ampicillin, lysing cells by sonication (20% power, 8 s), and plating the lysate on either Modified Bold Basal Media (MBBM; CCAP, UK), or modified artificial WC media (MWC; (59)) supplemented with amino acids, each with 1.5% agar and ampicillin. Plates were incubated at 25°C under 50 µE m^-2^s^-1^ light (10:14 L:D cycle). After 3—4 weeks, individual colonies were moved to standard algal growth conditions in liquid MBBM (25 °C, 50 µE m⁻² s⁻¹ light, 14:10 L:D cycle, 110 rpm shaking). Strain identity was confirmed by sequencing 18s rRNA and ITS2 regions (60).

### Selection in different nitrogen sources

In preparation for the selection experiment, fresh isolates were obtained as above. Six colonies were picked from each as founders for each biological replicate and moved to standard algal growth conditions in liquid MBBM. Each culture was pre-adapted for 3 days for their respective treatments, before being initiated at ∼1.2 × 10D cells mL⁻¹ in BBM media with one of three nitrogen sources, each providing 0.0088 M nitrogen: BBM-NO₃ (nitrate), BBM-ARG (arginine), or BBM-GLN (glutamine).

Populations were serially transferred weekly for 30 weeks by subculturing 25% of the total volume. Cell density was measured before and after each transfer using flow cytometry (CytoFLEX S, Beckman Coulter).

### Nitrogen use assay

To test nitrogen use of the evolved strains, the cultures were spun down (2000 rcf, 5 min), washed in BMM buffer (MBBM lacking sodium nitrate and peptone), and resuspended in 20 mL MBBM under standard growth conditions for 9 days to increase biomass. 10 mL of each culture was then pelleted and washed twice in BBM-buffer (final wash at 10,000 xg, 3 min). Pellets from each symbiont strain and selection treatment were then resuspended in either BBM-NO₃, BBM-ARG, or BBM-GLN, adjusted to OD₇₅₀ = 0.1, and transferred into 96-well plates in three technical replicates. OD₇₅₀ was measured at day 0 and day 7 using a plate reader (CLARIOstar Plus, BMG LABTECH).

### Symbiotic capacity assay

A common garden host, *P. bursaria* lab strain HA1, was used to test the symbiotic capacity of the evolved strains. HA1 cells were cured of their native symbionts by incubating in NCL with 20 µg mL^-1^ paraquat and 20 µg mL^-1^ cycloheximide in high light (50 µE m⁻² s⁻¹ light, 14:10 L:D cycle, 25 °C) for 12 days followed by 3–5 days in darkness and 5–7 days recovery in standard *Paramecium* conditions. Cells were inspected by microscopy to confirm the absence of symbionts and then reinfected with the evolved algal strains.

For reinfection, evolved algal strains were precultured for 14 days in MBBM, pelleted, and acclimated in NCL medium for 2 days. Symbiont-free HA1 hosts (10–15 cells) were then cocultured with ∼1.5 × 10D algal cells in 1.5 mL NCL medium under standard *Paramecium* conditions. Four control cultures without algae confirmed that hosts did not reestablish symbiosis with surviving native symbionts. After 14 days, reinfected HA1 were transferred to NCL supplemented with bacterial food (*Serratia marcescens*) and acclimated for 5 days before experimental assays.

The fitness effect for the HA1 hosts was quantified by directly competing the reinfected hosts with symbiont-free hosts in darkness (< 3 µE m⁻² s⁻¹) and high light (50 µE m⁻² s⁻¹). 10 cells of each were cultured in 1.5 mL of NCL (supplemented with *S. marcescens* on days 0 and 3) for 7 days. The final number of symbiotic and symbiont-free cells were quantified with flow cytometry (CytoFLEX S, Beckman Coulter).

### Sugar release assay

To quantify maltose and glucose secretion of evolved Symb-186b lines were cultured in either BBM-NO₃, BBM-ARG, or BBM-GLN without sucrose in standard growth conditions on a shaker (120 rpm) for 11 days to ensure cells were in exponential growth. Algal cultures were pelleted (2000 rcf, 15 min), washed with BBM-buffer and pelleted again (2000 rcf, 10 min). Three replicate 2 ml populations of 1.3 × 10^8^ cells mL^-1^ of each strain were resuspended (at equal concentration verified with flow cytometry) in pH 5.5 MBBM medium without N or sucrose (pH adjusted with 100 mM MES monohydrate and MES Na salt buffers) and incubated for 6 hours at standard growth conditions. After incubation all samples were filtered (0.22 µm) and the supernatant frozen (–20°C) for analysis with ion chromatography.

The sugar release samples were analysed with DIONEX ICS-6000 SP, AS-AP ion chromatography at Manchester Institute of Biotechnology (MIB, Mass Spectrometry & Separations Facility). The amount of maltose and glucose was determined using a CarboPac PA20 Guard Column (30 mm) and a CarboPac PA20 Analytical Column (150 mm). Multistep gradients of 45 min were created to optimise peak separation of both sugars, using 30 mM NaOH as the mobile phase with a sample injection volume of 10 µl. Sugar concentrations were then quantified using 6 concentrations (STD1, 5, 10, 50, 100, 250 ppm) of known standards.

### Phenotype analysis: phenotypes at transfer, nitrogen use and sugar release

All below analyses were carried out using R (v. 4.4.1; (61)), unless otherwise specified. Mixed-effects models were fit with `lme4::lmer()`(v. 1.1.37; (62)) to characterise each evolved strain’s phenotype at each transfer as well as sugar release and nitrogen use at the experiment end point. For cell density, per cell chlorophyll fluorescence, cell size and cell granularity at transfer, raw flow cytometry data was imported into R using `flowCore` (v. 2.16.0; (63)) and gated using the FITC.H channel (> 1000) to ensure only live cells were included. log(cells/ml), PerCP.A (chlorophyll fluorescence), FSC.A (forward scatter, cell size) and SSC.A (side scatter, cell granularity) were fit as responses with transfer number, symbiont and treatment as the fixed effects. For sugar release, maltose and glucose concentrations in ppm were fit as separate response variables with treatment as the fixed effect. For growth rate in different growth media, growth rate was calculated as log(OD750 at day 7/OD750 at day 0)/7 days and fit as the response variable with treatment, test medium and symbiont as the fixed effects. In all three cases, biological replicate was fit as a random effect with a variable intercept. Significance testing was done using `car::Anova()` (v. 3.1.3; (64)), and post hoc testing of marginal means as well as slopes and intercepts was carried out using the `emmeans` package (v.1.11.2; (65)).

### Phenotype analysis: symbiotic capacity

To obtain cell counts, the competition assay flow cytometry data was gated based on forward side scatter (i.e. size) to identify host cells. Reinfected cells were distinguished from symbiont-free cells based on single cell chlorophyll fluorescence (excitation 488 nm, emission 690 nm).

As a considerable proportion of the competition assay trials saw reinfected cells completely outcompeted by symbiont-free cells, fixation probability of the symbiont-free phenotype was tested by fitting a Bayesian generalised linear model (`brms` package, v. 2.22.0; (66)) with Bernoulli likelihood (logit link). Treatment, symbiont and light level were fit as fixed effects with weakly informative Normal (0,2) priors, while biological replicate was fit as a random effect with a variable intercept.

### DNA extraction and sequencing

Samples were collected for DNA sequencing from the 186b selection lines at transfers 1 and 30. Prior to extraction, 5 mL culture was grown in 25 mL MBBM for 14 days, pelleted (30,000 xg, 10 min), and flash-frozen in liquid nitrogen.

DNA was extracted using a custom CTAB–ethanol precipitation method. Cells were homogenised with 0.5 mm zirconium oxide beads in CTAB buffer, incubated with with 50 µl proteinase K (65 °C, 30 min), followed by addition of 3 µl RNase A and further incubation (37°C, 30 min), and then extracted with an 25:1 chloroform:isoamyl alcohol solution. DNA was pelleted (10,000 rpm, 60 min, 4 °C), washed twice with icecold 70% ethanol, and eluted in 200 µL TE buffer. Sequencing was carried out by CGR using the Illumina NovaSeq 6000 platform, using TruSeq PCR-free kit using a single lane of a S4 flow cell.

### Variant Calling and Allele Frequency Estimation

Paired-end Illumina reads were adapter and quality trimmed using Trim Galore! (v0.6; (67)) using standard settings. Following this, trimmed reads for each biological replicate (and including the ancestral library) were aligned to the Micractinium conductrix 186b nuclear genome assembly (19) using minimap2 (v2.28; (68)), with secondary alignments suppressed. Read group information indicating growing media condition and replicate number was added during alignment using SAMtools (v1.21; (69)). The next step included PCR and optical duplicate removal using GATK MarkDuplicates (v4.6.1; (70)), with duplicates retained in the output but marked to allow downstream filtering.

Variant discovery was performed using the GATK HaplotypeCaller in Genomic Variant Call Format (GVCF) mode (this allows records for all sites), with ploidy set to 1 (to reflect the haploid nature of the algal genome) and with one GVCF file per biological replicate. Then the individual GVCFs were combined per experimental condition using GATK CombineGVCFs, continuing with the ancestral sample being treated as an additional condition.

Joint genotyping was performed individually using GATK GenotypeGVCFs using the combined GCVF per condition and the reference genome. Finally, variant filtration was applied using GATK VariantFiltration; variant sites were filtered out if they showed low quality by depth (QD < 2.0), excessive strand bias by either Fisher Strand test or Strand Odds Ratio (FS > 60.0 or SOR > 4.0), poor mean mapping quality (MQ < 40.0), or insufficient read depth (DP < 10). The final passing variants were extracted using BCFtools (v1.21; Danecek et al 2021), and then per-sample genotype (GT) and allele depth (AD) fields were exported to a TSV file using bcftools query.

### Allele Frequency Calculation and Ancestral Filtering

Analyses for this section were performed in R (v4.4; (61)) using the tidyverse packages (71). The per-sample allele depth tables were reshaped from wide to long format, and allele frequencies (AF) were computed for each variant-sample combination as the ratio of ALT-supporting reads to total read depth. Sites with fewer than 10 total reads were excluded. To remove potential genetic variation present prior to experimental evolution, any variant position at which the ancestral sample exhibited an ALT allele frequency ≥ 0.1 was identified and then discarded from all evolved condition datasets.

### Functional Annotation

Variants passing ancestral filtering were functionally annotated using snpEff (v 5.2; (72)), with a custom database constructed from the *M. conductrix* 186b genome assembly, CDS sequences, protein sequences, and GFF3 gene models. The ANN field from the snpEff VCF output was parsed to recover effect class, predicted impact, gene name, gene identifier, and HGVS nomenclature for each alternate allele. Intergenic variants were excluded from down-stream analyses.

GO term mappings were derived from InterProScan (v 5.73; (73)) output, run against all available member databases with GO term and pathway lookup enabled. GO terms were extracted from the InterProScan TSV output and collapsed to a unique set per gene identi-fier.

### Genomic parallelism and candidate gene identification

Initial analyses were focussed on fixed or near-fixed variants in the peak of frequencies be-tween 0.95 and 1 (Fig. S7), with parallelism first assessed across the full dataset using these high-frequency variants. Presence–absence matrices were constructed and Jaccard dis-tances were calculated using the `vegan` package. Genome wide parallelism was tested by comparing within-versus between-treatment distances using permutation tests (n = 9999) and bootstrap resampling.

To identify candidate loci underlying treatment-specific divergence, the data set was ana-lysed separately for each treatment using multiple complementary approaches. First, Fisher’s exact tests were applied to identify individual SNPs or genes disproportionately as-sociated with one treatment over the others. Second, constrained ordinations (CAP) were performed using `vegan::capscale()`, with alleles/genes fitted as explanatory vectors via `envfit()` (v. 2.7.1; (74)); loci with significant loadings were ranked by effect size (r²). Third, an indicator species analysis was conducted using the `indicspecies` package (v. 1.8.0; (75)) to detect sets of loci indicative of a treatment. This was done across a range of allele frequency thresholds (1, 0.99, 0.97, 0.95, 0.90, 0.85, 0.80, 0.75) for determining fixed variants to ensure the results were robust to threshold changes. The output from these models was joined and filtered for variants unique to each treatment and parallel in at least 2 replicate populations and present in at least 75% of runs across the allele frequency thresholds to form our list of genes of interest.

### Candidate gene function identification

To summarise functional annotations, shortlisted genes were linked to Gene Ontology (GO) terms and analysed for GO enrichment against the full list of GO terms from the reference annotation using the `topGO` package (v. 2.56.0; (76)). Semantic similarity of enriched GO terms was calculated using the `rrvgo` package (v. 1.16.0; (77)) for plotting. Candidate protein sequences were further annotated by mapping to KEGG orthologs using GhostKOALA (https://www.kegg.jp/ghostkoala/) and by sequence similarity searches against the SwissProt Viridiplantae database using blastp through the NCBI web tool (https://blast.ncbi.nlm.nih.gov). IDs for the blastp output were mapped using the Uniprot web tool (https://www.uniprot.org/id-mapping).

### Untargeted metabolic fingerprinting using DESI-MSI

Culture samples were pelleted by centrifuging (2000 rcf, 5 min), weighed and flash frozen at transfers 1, 5, 10, 20 and 30 and stored in –80°C. Samples were dissolved in 1 mL of a methanol-water mix (4:1) for 10 minutes before being centrifuged (10 mins, 16 163 xg) and the supernatant removed. The supernatant was put in a SpeedVac (Thermo Fisher, UK) for 3 hours 30 minutes, until samples were completely dry. Samples were then reconstituted with 80:20 methanol water solvent at a ratio of 10 µL solvent per 1 mg wet mass.

2 µl of each sample was spotted onto Waters PTFE coated microscope glass slides (Waters Corporation, Wilmslow, UK) in a randomised block design (3 reps) and were analysed on a SYNAPT XS mass spectrometer (Waters Corporation, Wilmslow, UK) coupled with a modified DESI XS source in negative ionisation mode. DESI solvent spray was composed of a methanol-water mixture (98:2) at a flow rate of 1.5 μL/min with nitrogen gas flow set to 0.5—1 bar, a capillary voltage of 0.57 kV with 25 V sampling cone, heated transfer line at 450 °C and source temperature of 100 °C; the trap and transfer cell voltages were set at 4 and 2 V, respectively. We used a resolution of 150 µm moving at 1500 µm/s. The mass spectrometer was operated using MassLynx v4.2 (Waters Corporation, Wilmslow, UK). Deposited spot regions were extracted and pre-processed (0.2 Da window, m/z between 50-2000 were selected, top 2000 peaks) using HDI software (v1.5). Data were further processed using R for lock mass correction, clustering of peaks with a 10 ppm tolerance and calculating median weighted centroids, requiring presence in at least 2/3 technical replicates and keeping only features above a minimum intensity threshold (sample mean > median of all means in sample) to reduce downstream dimensionality.

### Identification of candidate features

To test the distribution of *m/z* separated features, the *m/z* intensity matrix was log + 1 transformed as well as Pareto scaled and a PCA was fit using vegan::rda() (v. 2.7.1; (74)). To determine whether clustering was explained by treatment, a PERMANOVA was fit using vegan::adonis2() (v. 2.7.1; (74)).

To identify *m/z* features of interest, PCAs were performed on the full dataset as well as sub-sets comprising a single treatment or a single timepoint and the top 100 loadings were extracted. Furthermore, as a complementary approach, further m/z peaks of interest were identified by using a random forest model fit with `ranger` (v. 0.17.0; (78)) using 70% of the data frame at each permutation to allow for estimating permutation importance of each feature (n = 1000), selecting features present in the in top 2x of the elbow of the mean importance curve in at least 50% of all permutations. An LMM (intensity ∼ treatment * transfer + (1|bio_rep)) was fit to each m/z of interest `lme4::lmer()`(v. 1.1.37; (62)), and m/z peaks where treatment or the interaction with transfer had a significant effect were shortlisted. The shortlist was further filtered based on there being a significant difference as well as a minimum 1.5 fold change between at least two of the treatments by the final transfer.

### MS/MS methods

The top features from the statistical analysis were targeted for DESI-MS/MS analysis. Data was acquired using MassLynx V4.2 (Waters Corporation, Wilmslow, UK) by rastering over the sample slides. Trap collision voltage was optimised for each quadruple isolated *m/z* value, ranging from 4-45 V, with all other MS conditions maintained as those used during the full scan DESI-MS experiment. Putative ID’s were assigned using product ion *m/z* and MS/MS fragment, comparing to literature (79–83), the LIPIDMAPS and HMBD database.

Characteristic headgroup fragments were used to confirm lipid class, *m/z* 225 [C_6_H_9_O_7_S]^−^ for sulfonquinovosyl diagcylglycerol (SQDG) (79), *m/z* 241 [C_6_H_10_O_8_P]^-^ for phosphatidylinositol (PI) and *m/z* 153 [C_3_H_6_O_5_P]^-^ for all glycerophospholipids (84). Additionally, fatty acyl chain fragments and neutral loss were used for further confirmation. MS/MS data for features that could not be assigned an identity can be found in Supplementary Materials 3.

