## Supplementary materials 1 for "Adaptation to free-living drives loss of beneficial endosymbiosis through metabolic trade-offs"

**Table S1 –** Test statistics from mixed effects model of effect of selection treatment, transfer number and symbiont on day 7 cell density along with estimated trends.

| **Cell density: Mixed effects model** | | | | | |
| --- | --- | --- | --- | --- | --- |
| *Main effect* | | | *Chisq* | *DF* | *p* |
| Transfer | | | 453.6 | 1 | < 0.001 |
| Symbiont | | | 2612.8 | 3 | < 0.001 |
| Treatment | | | 7892.6 | 2 | < 0.001 |
| Transfer:Symbiont | | | 180.3 | 3 | < 0.001 |
| Transfer:Treatment | | | 20.2 | 2 | < 0.001 |
| Symbiont:Treatment | | | 5878.5 | 6 | < 0.001 |
| Transfer:Symbiont:Treatment | | | 211.9 | 6 | < 0.001 |
| **Cell density: Estimated trends** | | | | | |
| *Symbiont* | *Treatment* | *Transfer trend* | *DF* | *Lower CL* | *Upper CL* |
| 186b | A | 0.025 | 2125 | 0.019 | 0.031 |
|  | G | 0.041 | 2125 | 0.035 | 0.047 |
|  | N | 0.020 | 2125 | 0.014 | 0.026 |
| 21 | A | 0.019 | 2127 | 0.012 | 0.025 |
|  | G | 0.008 | 2125 | 0.002 | 0.014 |
|  | N | -0.023 | 2125 | -0.029 | -0.018 |
| 37 | A | 0.020 | 2125 | 0.014 | 0.026 |
|  | G | 0.016 | 2125 | 0.010 | 0.022 |
|  | N | 0.044 | 2125 | 0.039 | 0.051 |
| HZ75.5 | A | 0.034 | 2125 | 0.028 | 0.040 |
|  | G | 0.006 | 2125 | 0.0001 | 0.012 |
|  | N | 0.020 | 2125 | 0.014 | 0.026 |
| **Mean fluorescence: Mixed effects model** | | | | | |
| *Main effect* | | | *Chisq* | *DF* | *p* |
| Transfer | | | 673.5 | 1 | < 0.001 |
| Symbiont | | | 3845.7 | 3 | < 0.001 |
| Treatment | | | 2638.3 | 2 | < 0.001 |
| Transfer:Symbiont | | | 83.8 | 3 | < 0.001 |
| Transfer:Treatment | | | 5.4 | 2 | 0.07 |
| Symbiont:Treatment | | | 4773.0 | 6 | < 0.001 |
| Transfer:Symbiont:Treatment | | | 52.1 | 6 | < 0.001 |
| **Mean fluorescence: Estimated trends** | | | | | |
| *Symbiont* | *Treatment* | *Transfer trend* | *DF* | *Lower CL* | *Upper CL* |
| 186b | A | 0.023 | 2125 | 0.018 | 0.027 |
|  | G | 0.036 | 2125 | 0.031 | 0.040 |
|  | N | 0.019 | 2125 | 0.015 | 0.024 |
| 21 | A | 0.019 | 2126 | 0.014 | 0.023 |
|  | G | 0.013 | 2125 | 0.008 | 0.017 |
|  | N | 0.008 | 2125 | 0.004 | 0.012 |
| 37 | A | 0.006 | 2125 | 0.002 | 0.011 |
|  | G | 0.008 | 2125 | 0.003 | 0.012 |
|  | N | 0.017 | 2125 | 0.013 | 0.022 |
| HZ75.5 | A | 0.019 | 2125 | 0.015 | 0.024 |
|  | G | 0.018 | 2125 | 0.013 | 0.022 |
|  | N | 0.015 | 2125 | 0.011 | 0.020 |
| **Cell size: Mixed effects model** | | | | | |
| *Main effect* | | | *Chisq* | *DF* | *p* |
| Transfer | | | 236.7 | 1 | < 0.001 |
| Symbiont | | | 1109.5 | 3 | < 0.001 |
| Treatment | | | 1086.1 | 2 | < 0.001 |
| Transfer:Symbiont | | | 199.1 | 3 | < 0.001 |
| Transfer:Treatment | | | 109.9 | 2 | < 0.001 |
| Symbiont:Treatment | | | 1208.7 | 6 | < 0.001 |
| Transfer:Symbiont:Treatment | | | 215.1 | 6 | < 0.001 |
| **Cell size: Estimated trends** | | | | | |
| *Symbiont* | *Treatment* | *Transfer trend* | *DF* | *Lower CL* | *Upper CL* |
| 186b | A | 0.001 | 2125 | -0.0008 | 0.003 |
|  | G | 0.015 | 2125 | 0.013 | 0.017 |
|  | N | 0.008 | 2125 | 0.006 | 0.010 |
| 21 | A | 0.005 | 2126 | 0.003 | 0.007 |
|  | G | 0.006 | 2125 | 0.004 | 0.008 |
|  | N | -0.010 | 2125 | -0.012 | -0.008 |
| 37 | A | -0.0003 | 2125 | -0.002 | 0.001 |
|  | G | 0.002 | 2125 | 0.0003 | 0.004 |
|  | N | 0.002 | 2125 | 0.0004 | 0.004 |
| HZ75.5 | A | 0.006 | 2125 | 0.005 | 0.008 |
|  | G | 0.009 | 2125 | 0.007 | 0.011 |
|  | N | 0.008 | 2125 | 0.006 | 0.010 |
| **Cell granularity: Mixed effects model** | | | | | |
| *Main effect* | | | *Chisq* | *DF* | *p* |
| Transfer | | | 0.03 | 1 | 0.85 |
| Symbiont | | | 942.2 | 3 | < 0.001 |
| Treatment | | | 227.4 | 2 | < 0.001 |
| Transfer:Symbiont | | | 239.2 | 3 | < 0.001 |
| Transfer:Treatment | | | 177.8 | 2 | < 0.001 |
| Symbiont:Treatment | | | 528.8 | 6 | < 0.001 |
| Transfer:Symbiont:Treatment | | | 301.9 | 6 | < 0.001 |
| **Cell granularity: Estimated trends** | | | | | |
| *Symbiont* | *Treatment* | *Transfer trend* | *DF* | *Lower CL* | *Upper CL* |
| 186b | A | -0.016 | 2125 | -0.019 | -0.013 |
|  | G | 0.011 | 2125 | 0.008 | 0.014 |
|  | N | 0.004 | 2125 | 0.002 | 0.007 |
| 21 | A | -0.0002 | 2126 | -0.003 | 0.003 |
|  | G | 0.008 | 2125 | 0.005 | 0.019 |
|  | N | -0.017 | 2125 | -0.019 | -0.014 |
| 37 | A | -0.004 | 2125 | -0.007 | -0.002 |
|  | G | 0.001 | 2125 | -0.002 | 0.004 |
|  | N | 0.015 | 2125 | -0.017 | -0.012 |
| HZ75.5 | A | 0.004 | 2125 | 0.001 | 0.007 |
|  | G | 0.011 | 2125 | 0.008 | 0.014 |
|  | N | 0.017 | 2125 | 0.014 | 0.020 |

**Table S2 –** Test statistics from mixed effects model of effect of selection treatment, nitrogen medium, symbiont on growth rate change compared to the ancestor along with estimated marginal means with 95% confidence intervals.

| **Mixed effects model** | | | | | | | |
| --- | --- | --- | --- | --- | --- | --- | --- |
| *Main effect* | | | *Chisq* | | | *DF* | *p* |
| Selection treatment (evo_trt) | | | 4.5 | | | 2 | 0.1 |
| Nitrogen medium (test_trt) | | | 36.5 | | | 2 | < 0.001 |
| Symbiont | | | 54.1 | | | 3 | < 0.001 |
| Evo_trt:test_trt | | | 28.6 | | | 4 | < 0.001 |
| Evo_trt:symbiont | | | 37.9 | | | 6 | < 0.001 |
| Test_trt:symbiont | | | 70.6 | | | 6 | < 0.001 |
| Evo_trt:test_trt:symbiont | | | 32.9 | | | 12 | < 0.001 |
| **Estimated marginal means with 95% confidence intervals** | | | | | | | |
| *Symbiont* | *Selection treatment* | *Growth test medium* | *emmean* | *SE* | *df* | *Lower CL* | *Upper CL* |
| 186b | Arginine | Arginine | 0.023 | 0.014 | 177 | -0.005 | 0.051 |
|  | Glutamine |  | 0.023 | 0.014 | 177 | -0.005 | 0.051 |
|  | Nitrate |  | 0.034 | 0.014 | 177 | 0.006 | 0.062 |
|  | Arginine | Glutamine | 0.075 | 0.014 | 177 | 0.047 | 0.103 |
|  | Glutamine |  | 0.196 | 0.014 | 177 | 0.168 | 0.224 |
|  | Nitrate |  | 0.040 | 0.014 | 177 | 0.012 | 0.068 |
|  | Arginine | Nitrate | 0.032 | 0.014 | 177 | 0.004 | 0.060 |
|  | Glutamine |  | 0.010 | 0.014 | 177 | -0.018 | 0.038 |
|  | Nitrate |  | 0.024 | 0.014 | 177 | -0.004 | 0.052 |
| 21 | Arginine | Arginine | 0.006 | 0.016 | 177 | -0.025 | 0.036 |
|  | Glutamine |  | -0.012 | 0.014 | 177 | -0.040 | 0.016 |
|  | Nitrate |  | 0.034 | 0.014 | 177 | 0.006 | 0.062 |
|  | Arginine | Glutamine | 0.016 | 0.016 | 177 | -0.015 | 0.047 |
|  | Glutamine |  | 0.007 | 0.014 | 177 | -0.021 | 0.035 |
|  | Nitrate |  | 0.045 | 0.014 | 177 | 0.017 | 0.072 |
|  | Arginine | Nitrate | 0.016 | 0.016 | 177 | -0.015 | 0.047 |
|  | Glutamine |  | 0.017 | 0.014 | 177 | -0.011 | 0.045 |
|  | Nitrate |  | 0.059 | 0.014 | 177 | 0.031 | 0.087 |
| 37 | Arginine | Arginine | -0.010 | 0.014 | 177 | -0.038 | 0.017 |
|  | Glutamine |  | 0.005 | 0.014 | 177 | -0.023 | 0.033 |
|  | Nitrate |  | -0.010 | 0.014 | 177 | -0.038 | 0.018 |
|  | Arginine | Glutamine | 0.067 | 0.014 | 177 | 0.039 | 0.095 |
|  | Glutamine |  | 0.082 | 0.014 | 177 | 0.054 | 0.110 |
|  | Nitrate |  | 0.041 | 0.014 | 177 | 0.013 | 0.069 |
|  | Arginine | Nitrate | 0.040 | 0.014 | 177 | 0.012 | 0.068 |
|  | Glutamine |  | 0.014 | 0.014 | 177 | -0.014 | 0.042 |
|  | Nitrate |  | 0.032 | 0.014 | 177 | 0.004 | 0.060 |
| HZ75.5 | Arginine | Arginine | -0.002 | 0.014 | 177 | -0.030 | 0.026 |
|  | Glutamine |  | 0.002 | 0.014 | 177 | -0.026 | 0.030 |
|  | Nitrate |  | 0.035 | 0.014 | 177 | 0.008 | 0.063 |
|  | Arginine | Glutamine | -0.025 | 0.014 | 177 | -0.053 | 0.003 |
|  | Glutamine |  | -0.004 | 0.014 | 177 | -0.032 | 0.024 |
|  | Nitrate |  | -0.003 | 0.014 | 177 | -0.031 | 0.025 |
|  | Arginine | Nitrate | -0.012 | 0.014 | 177 | -0.040 | 0.016 |
|  | Glutamine |  | -0.005 | 0.014 | 177 | -0.033 | 0.023 |
|  | Nitrate |  | 0.037 | 0.014 | 177 | 0.009 | 0.065 |

**Table S3 –** Model summary output and selected odds ratio contrasts from Bayesian generalised linear model estimating probability of non-symbiont host phenotype fixing based on light level, selection treatment and strain. Model support for an effect or difference is strong when the confidence interval does not overlap 0 for an estimate or 1 for an odds ratio.

| **Model summary output** | | | | | |
| --- | --- | --- | --- | --- | --- |
| *Parameter* | | *Estimate* | *Est. Error* | *l-95% CI* | *u-95% CI* |
| Intercept | | -1.84 | 0.45 | -2.76 | -0.97 |
| treatmentG | | 0.36 | 0.57 | -0.78 | 1.49 |
| treatmentN | | 2.54 | 0.53 | 1.48 | 3.60 |
| light.L | | -1.44 | 0.55 | -2.56 | -0.42 |
| strain21 | | -0.32 | 0.69 | -1.76 | 0.99 |
| strain37 | | -0.37 | 0.65 | -1.65 | 0.88 |
| strainHZ75.5 | | -0.69 | 0.69 | -2.09 | 0.63 |
| treatmentG:light.L | | 0.42 | 0.73 | -0.97 | 1.87 |
| treatmentN:light.L | | 1.03 | 0.67 | -0.25 | 2.39 |
| treatmentG:strain21 | | -1.24 | 1.06 | -3.44 | 0.69 |
| treatmentN:strain21 | | -2.11 | 0.86 | -3.79 | -0.43 |
| treatmentG:strain37 | | 0.05 | 0.83 | -1.53 | 1.69 |
| treatmentN:strain37 | | -2.28 | 0.93 | -4.12 | -0.49 |
| treatmentG:strainHZ75.5 | | 0.99 | 0.83 | -0.62 | 2.66 |
| treatmentN:strainHZ75.5 | | -1.07 | 0.83 | -2.69 | 0.57 |
| light.L:strain21 | | -0.86 | 0.88 | -2.60 | 0.84 |
| light.L:strain37 | | -0.60 | 0.84 | -2.31 | 0.97 |
| light.L:strainHZ75.5 | | -0.95 | 0.88 | -2.69 | 0.69 |
| treatmentG:light.L:strain21 | | -0.96 | 0.30 | -3.66 | 1.55 |
| treatmentN:light.L:strain21 | | -0.24 | 1.10 | -2.40 | 1.90 |
| treatmentG:light.L:strain37 | | 0.81 | 1.06 | -1.22 | 2.89 |
| treatmentN:light.L:strain37 | | -1.75 | 1.20 | -4.11 | 0.52 |
| treatmentG:light.L:strainHZ75.5 | | 0.96 | 1.07 | -1.06 | 3.05 |
| treatmentN:light.L:strainHZ75.5 | | 0.41 | 1.05 | -1.62 | 2.44 |
| *Random effect:* bio.rep, sd(Intercept) | | 0.23 | 0.20 | 0.01 | 0.75 |
| **Selected odds ratio contrasts** | | | | | |
| *Symbiont* | *Light level* | *Contrast* | *Odds ratio* | *Lower HPD* | *Upper HPD* |
| 186b | Darkness | A / G | 0.94 | 0.17 | 2.61 |
| 186b | Darkness | A / N | 0.16 | 0.03 | 0.45 |
| 186b | Darkness | G / N | 0.18 | 0.02 | 0.54 |
| 186b | High light | A / G | 0.51 | 0.02 | 2.29 |
| 186b | High light | A / N | 0.04 | 0 | 0.13 |
| 186b | High light | G / N | 0.08 | 0 | 0.28 |
| 21 | Darkness | A / G | 1.63 | 0.18 | 5.53 |
| 21 | Darkness | A / N | 1.13 | 0.13 | 3.73 |
| 21 | Darkness | G / N | 0.7 | 0.06 | 2.49 |
| 21 | High light | A / G | 3.14 | 0.01 | 91.87 |
| 21 | High light | A / N | 0.39 | 0 | 3.2 |
| 21 | High light | G / N | 0.13 | 0 | 2.14 |
| 37 | Darkness | A / G | 1.56 | 0.14 | 5.54 |
| 37 | Darkness | A / N | 0.47 | 0.07 | 1.59 |
| 37 | Darkness | G / N | 0.29 | 0.01 | 1.01 |
| 37 | High light | A / G | 0.29 | 0 | 2.01 |
| 37 | High light | A / N | 1.2 | 0.01 | 20.72 |
| 37 | High light | G / N | 4.06 | 0.04 | 72.65 |
| HZ75.5 | Darkness | A / G | 0.69 | 0.09 | 2.16 |
| HZ75.5 | Darkness | A / N | 0.64 | 0.09 | 2.14 |
| HZ75.5 | Darkness | G / N | 0.93 | 0.12 | 3.17 |
| HZ75.5 | High light | A / G | 0.1 | 0 | 0.76 |
| HZ75.5 | High light | A / N | 0.09 | 0 | 0.59 |
| HZ75.5 | High light | G / N | 0.84 | 0.03 | 4.01 |

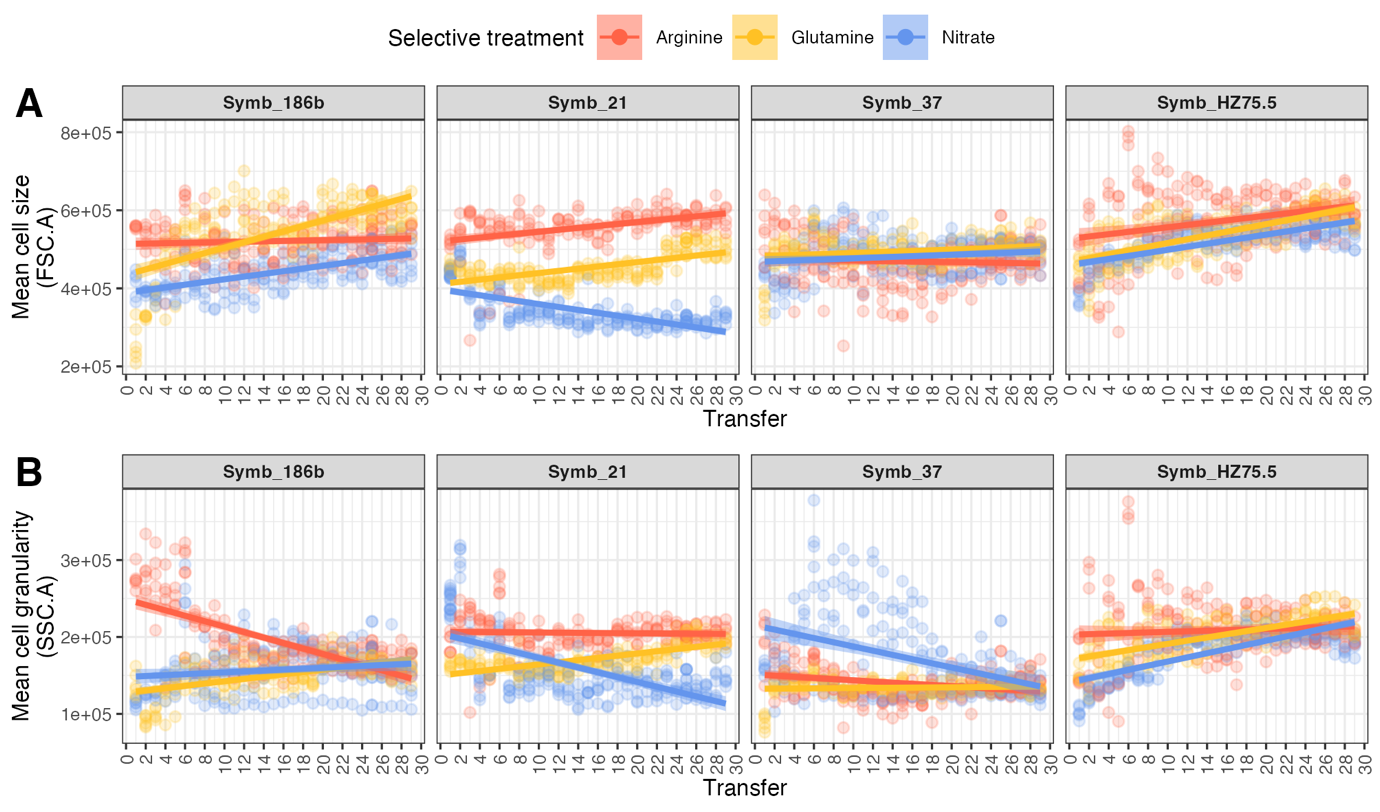

**Figure S1 – Cell size and cell granularity across transfers**

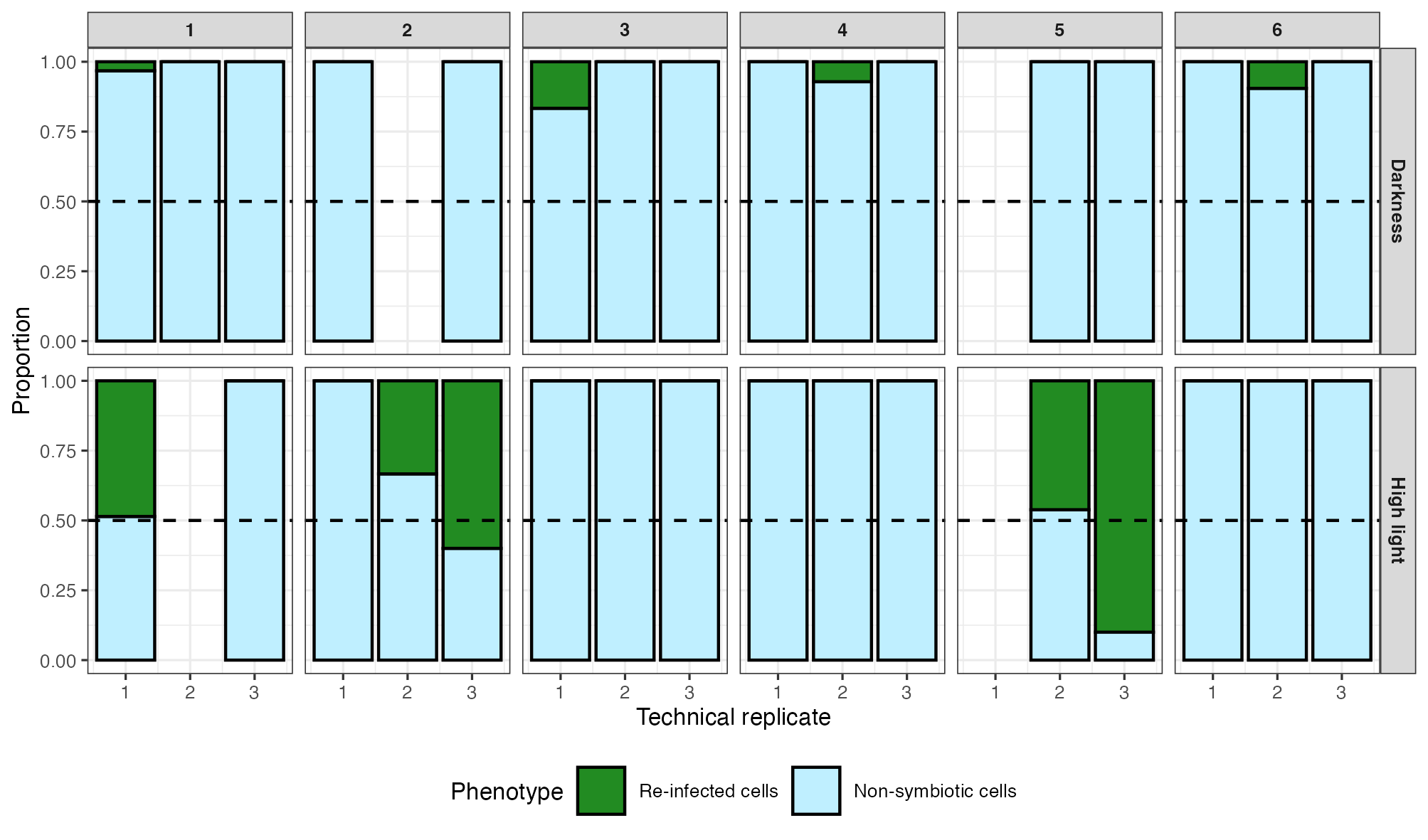

**Figure S2 – Proportion of non-symbiotic cells and host cells reinfected with nitrate-evolved Symb_186b.** The facet label numbers correspond to biological replicate.

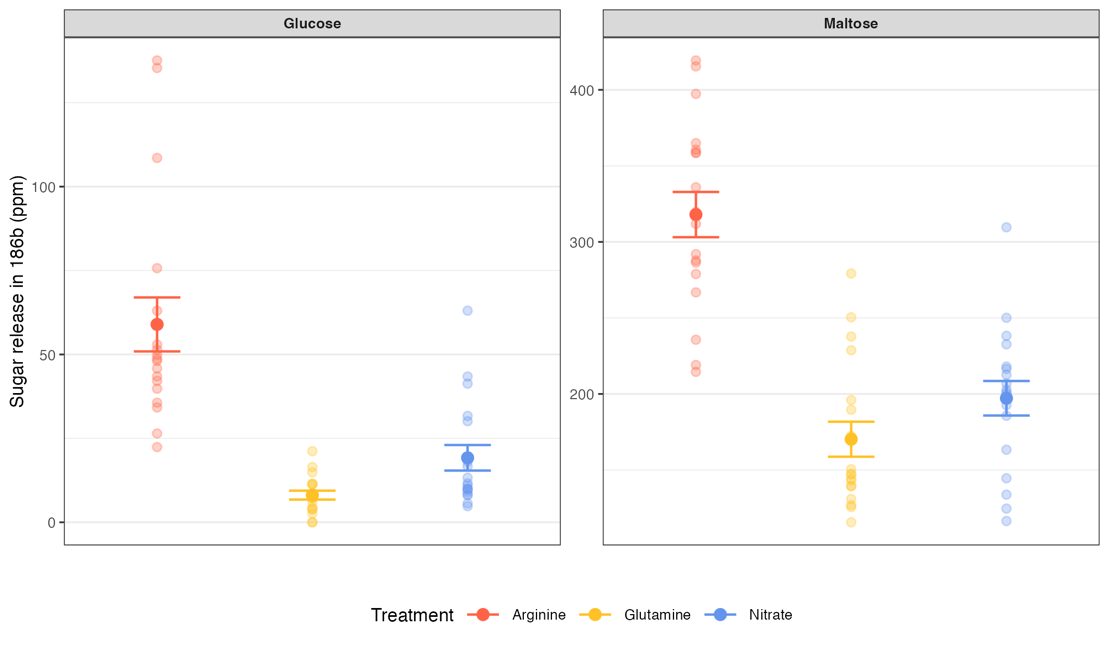

**Figure S3 – Maltose and glucose release in ppm by evolved Symb_186b algal strains**.  Raw data are shown as semi-transparent points with means and standard error represented as opaque points and error bars.

**Table S4 –** Test statistics from mixed effects model of effect of treatment on sugar release along with *post hoc* pairwise contrasts.

| **Mixed effects model** | | | | | **Pairwise contrasts** | | | |
| --- | --- | --- | --- | --- | --- | --- | --- | --- |
| *Sugar* | *Main effect* | *Chisq* | *DF* | *p* | *contrast* | *t-ratio* | *DF* | *p* |
| Maltose | Treatment | 77.9 | 2 | < 0.001 | A—G | 8.5 | 46 | < 0.001 |
|  |  |  |  |  | A—N | 6.5 | 46 | < 0.001 |
|  |  |  |  |  | G—N | -2.01 | 46 | 0.12 |
| Glucose | Treatment | 85.784 | 2 | < 0.001 | A—G | 9.2 | 46 | < 0.001 |
|  |  |  |  |  | A—N | 5.5 | 46 | < 0.001 |
|  |  |  |  |  | G—N | -3.7 | 46 | 0.002 |

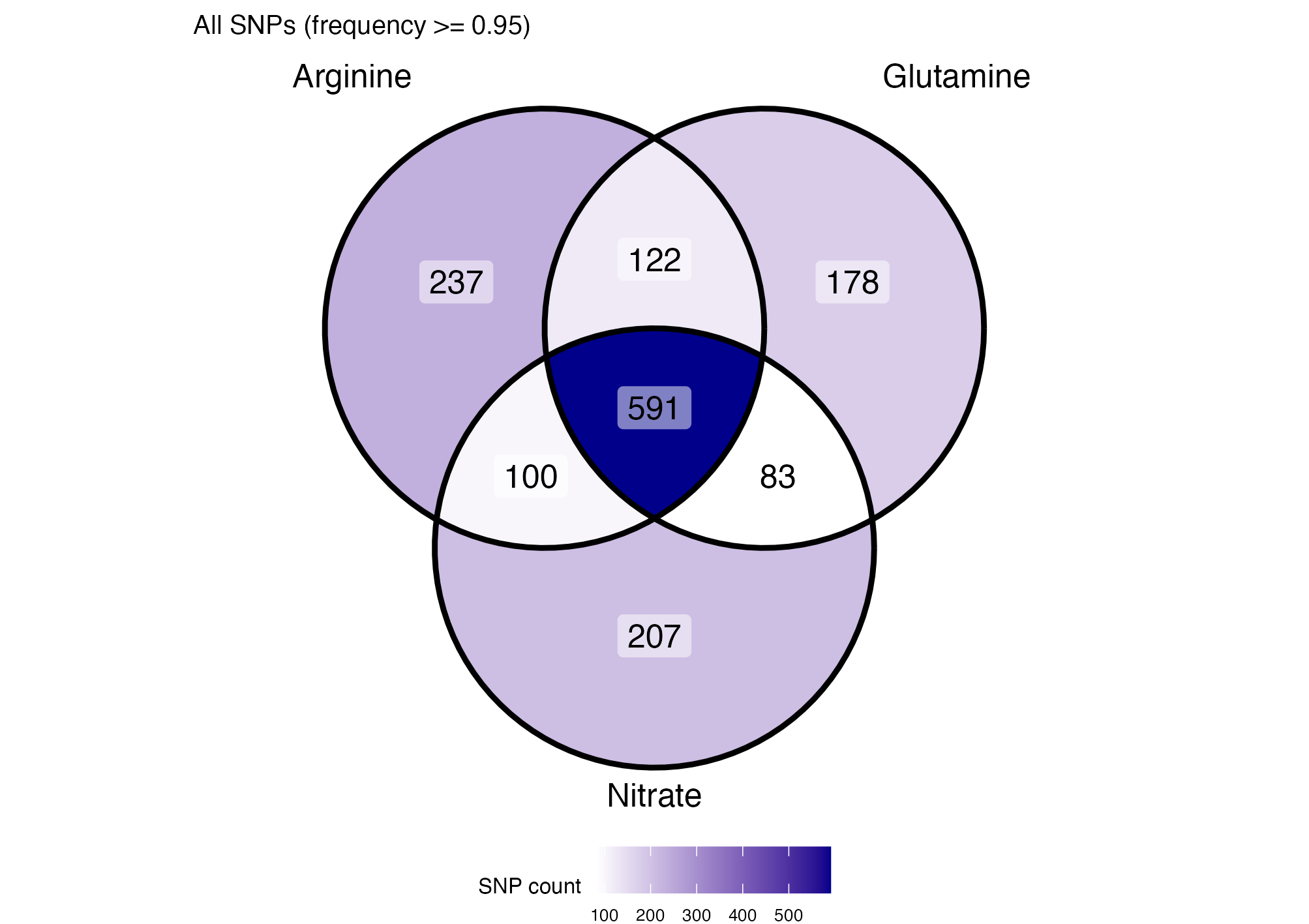

**Figure S4** –Distribution of high frequency (>= 0.95) SNPs among treatments.

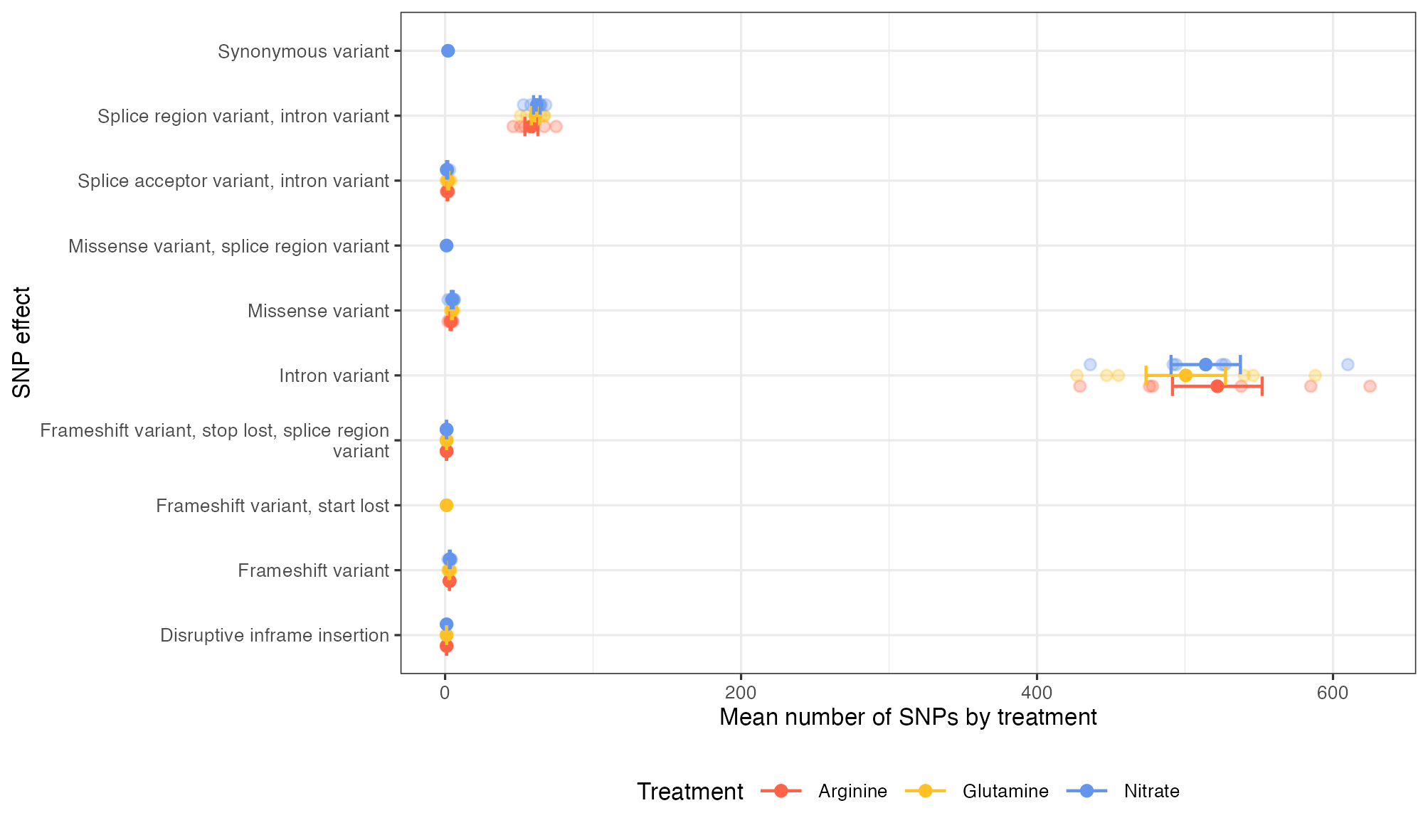

**Figure S5 – Distribution of SNP effects by treatment.** Raw data are shown as semi-transparent points with means and standard error represented as opaque points and error bars.

**Table S5 – GO enrichment analysis of shortlist genes against background of all genes with SNPs in the dataset. Only terms with p < 0.05 have been included.**

| **Treatment** | **Ontology** | **GO.ID** | **Term** | **Annotated** | **Significant** | **Expected** | **p** |
| --- | --- | --- | --- | --- | --- | --- | --- |
| **Arginine** | **BP** | GO:1901700 | response to oxygen-containing compound | 21 | 4 | 0.3 | 0.0002 |
|  |  | GO:0016043 | cellular component organization | 72 | 5 | 1.04 | 0.0034 |
|  |  | GO:0044281 | small molecule metabolic process | 55 | 4 | 0.79 | 0.0075 |
|  |  | GO:0007049 | cell cycle | 11 | 2 | 0.16 | 0.0103 |
|  |  | GO:0071422 | succinate transmembrane transport | 11 | 2 | 0.16 | 0.0103 |
|  |  | GO:0035433 | acetate transmembrane transport | 11 | 2 | 0.16 | 0.0103 |
|  |  | GO:0015711 | organic anion transport | 11 | 2 | 0.16 | 0.0103 |
|  |  | GO:0015718 | monocarboxylic acid transport | 11 | 2 | 0.16 | 0.0103 |
|  |  | GO:0015740 | C4-dicarboxylate transport | 11 | 2 | 0.16 | 0.0103 |
|  |  | GO:0015744 | succinate transport | 11 | 2 | 0.16 | 0.0103 |
|  |  | GO:0051726 | regulation of cell cycle | 11 | 2 | 0.16 | 0.0103 |
|  |  | GO:0006846 | acetate transport | 11 | 2 | 0.16 | 0.0103 |
|  |  | GO:0006835 | dicarboxylic acid transport | 11 | 2 | 0.16 | 0.0103 |
|  |  | GO:1903825 | organic acid transmembrane transport | 11 | 2 | 0.16 | 0.0103 |
|  |  | GO:1905039 | carboxylic acid transmembrane transport | 11 | 2 | 0.16 | 0.0103 |
|  |  | GO:0015849 | organic acid transport | 11 | 2 | 0.16 | 0.0103 |
|  |  | GO:0046942 | carboxylic acid transport | 11 | 2 | 0.16 | 0.0103 |
|  |  | GO:0031324 | negative regulation of cellular metabolic process | 13 | 2 | 0.19 | 0.0143 |
|  |  | GO:0031327 | negative regulation of cellular biosynthetic | 13 | 2 | 0.19 | 0.0143 |
|  |  | GO:1902679 | negative regulation of RNA biosynthetic process | 13 | 2 | 0.19 | 0.0143 |
|  |  | GO:0009890 | negative regulation of biosynthetic process | 13 | 2 | 0.19 | 0.0143 |
|  |  | GO:0009892 | negative regulation of metabolic process | 13 | 2 | 0.19 | 0.0143 |
|  |  | GO:0045892 | negative regulation of DNA-templated transcription | 13 | 2 | 0.19 | 0.0143 |
|  |  | GO:0048523 | negative regulation of cellular process | 13 | 2 | 0.19 | 0.0143 |
|  |  | GO:0051172 | negative regulation of nitrogen compound... | 13 | 2 | 0.19 | 0.0143 |
|  |  | GO:0048519 | negative regulation of biological process | 13 | 2 | 0.19 | 0.0143 |
|  |  | GO:0045934 | negative regulation of nucleobase-containing compound metabolic process | 13 | 2 | 0.19 | 0.0143 |
|  |  | GO:0051253 | negative regulation of RNA metabolic process | 13 | 2 | 0.19 | 0.0143 |
|  |  | GO:0010558 | negative regulation of macromolecule biosynthetic process | 13 | 2 | 0.19 | 0.0143 |
|  |  | GO:0010605 | negative regulation of macromolecule metabolic process | 13 | 2 | 0.19 | 0.0143 |
|  |  | GO:0006400 | tRNA modification | 39 | 3 | 0.56 | 0.0178 |
|  |  | GO:0010033 | response to organic substance | 43 | 3 | 0.62 | 0.0231 |
|  |  | GO:0018193 | peptidyl-amino acid modification | 20 | 2 | 0.29 | 0.0327 |
|  |  | GO:0170033 | L-amino acid metabolic process | 20 | 2 | 0.29 | 0.0327 |
|  |  | GO:0170039 | proteinogenic amino acid metabolic process | 20 | 2 | 0.29 | 0.0327 |
|  |  | GO:1901605 | alpha-amino acid metabolic process | 22 | 2 | 0.32 | 0.0391 |
|  |  | GO:0009451 | RNA modification | 56 | 3 | 0.81 | 0.0456 |
|  | **MF** | GO:0031593 | polyubiquitin modification-dependent protein binding | 11 | 2 | 0.09 | 0.0035 |
|  |  | GO:0015360 | acetate:proton symporter activity | 11 | 2 | 0.09 | 0.0035 |
|  |  | GO:0003714 | transcription corepressor activity | 17 | 2 | 0.14 | 0.0083 |
|  |  | GO:0000981 | DNA-binding transcription factor activity, RNA polymerase II-specific | 23 | 2 | 0.19 | 0.0150 |
|  |  | GO:0016788 | hydrolase activity, acting on ester bonds | 23 | 2 | 0.19 | 0.0150 |
|  |  | GO:0016874 | ligase activity | 68 | 3 | 0.56 | 0.0180 |
|  | **CC** | GO:0032991 | protein-containing complex | 225 | 9 | 1.83 | 0.0007 |
|  |  | GO:0005667 | transcription regulator complex | 15 | 2 | 0.12 | 0.0063 |
|  |  | GO:0098796 | membrane protein complex | 32 | 2 | 0.26 | 0.0273 |
| **Glutamine** | **BP** | GO:0000302 | response to reactive oxygen species | 14 | 3 | 0.25 | 0.0017 |
|  |  | GO:0042744 | hydrogen peroxide catabolic process | 15 | 3 | 0.27 | 0.0021 |
|  |  | GO:0051168 | nuclear export | 17 | 3 | 0.3 | 0.0030 |
|  |  | GO:1901605 | alpha-amino acid metabolic process | 22 | 3 | 0.39 | 0.0064 |
|  |  | GO:0006357 | regulation of transcription by RNA polymerase II | 69 | 5 | 1.23 | 0.0070 |
|  |  | GO:0070887 | cellular response to chemical stimulus | 49 | 4 | 0.87 | 0.0104 |
|  |  | GO:0034599 | cellular response to oxidative stress | 27 | 3 | 0.48 | 0.0115 |
|  |  | GO:0062197 | cellular response to chemical stress | 27 | 3 | 0.48 | 0.0115 |
|  |  | GO:0007049 | cell cycle | 11 | 2 | 0.2 | 0.0154 |
|  |  | GO:0051726 | regulation of cell cycle | 11 | 2 | 0.2 | 0.0154 |
|  |  | GO:0000027 | ribosomal large subunit assembly | 13 | 2 | 0.23 | 0.0213 |
|  |  | GO:0042273 | ribosomal large subunit biogenesis | 13 | 2 | 0.23 | 0.0213 |
|  |  | GO:0070925 | organelle assembly | 16 | 2 | 0.28 | 0.0317 |
|  |  | GO:0042255 | ribosome assembly | 16 | 2 | 0.28 | 0.0317 |
|  |  | GO:0140694 | non-membrane-bounded organelle assembly | 16 | 2 | 0.28 | 0.0317 |
|  |  | GO:0170034 | L-amino acid biosynthetic process | 17 | 2 | 0.3 | 0.0355 |
|  |  | GO:0170038 | proteinogenic amino acid biosynthetic process | 17 | 2 | 0.3 | 0.0355 |
|  |  | GO:1901607 | alpha-amino acid biosynthetic process | 17 | 2 | 0.3 | 0.0355 |
|  |  | GO:0071826 | protein-RNA complex organization | 18 | 2 | 0.32 | 0.0395 |
|  |  | GO:0022618 | protein-RNA complex assembly | 18 | 2 | 0.32 | 0.0395 |
|  |  | GO:0065003 | protein-containing complex assembly | 18 | 2 | 0.32 | 0.0395 |
|  |  | GO:0008652 | amino acid biosynthetic process | 20 | 2 | 0.36 | 0.0480 |
|  |  | GO:0044283 | small molecule biosynthetic process | 20 | 2 | 0.36 | 0.0480 |
|  |  | GO:0046394 | carboxylic acid biosynthetic process | 20 | 2 | 0.36 | 0.0480 |
|  |  | GO:0170033 | L-amino acid metabolic process | 20 | 2 | 0.36 | 0.0480 |
|  |  | GO:0170039 | proteinogenic amino acid metabolic process | 20 | 2 | 0.36 | 0.0480 |
|  |  | GO:0016053 | organic acid biosynthetic process | 20 | 2 | 0.36 | 0.0480 |
|  | **MF** | GO:0004601 | peroxidase activity | 22 | 3 | 0.24 | 0.0016 |
|  |  | GO:0016740 | transferase activity | 888 | 18 | 9.52 | 0.0042 |
|  |  | GO:0004252 | serine-type endopeptidase activity | 155 | 6 | 1.66 | 0.0060 |
|  |  | GO:0140098 | catalytic activity, acting on RNA | 117 | 5 | 1.25 | 0.0080 |
|  |  | GO:0016772 | transferase activity, transferring phosphorus containing groups | 343 | 9 | 3.68 | 0.0101 |
|  |  | GO:0140096 | catalytic activity, acting on a protein | 757 | 18 | 8.12 | 0.0132 |
|  |  | GO:0004672 | protein kinase activity | 316 | 8 | 3.39 | 0.0186 |
|  |  | GO:0042910 | xenobiotic transmembrane transporter activity | 20 | 2 | 0.21 | 0.0190 |
|  |  | GO:0016773 | phosphotransferase activity, alcohol groups as acceptor | 318 | 8 | 3.41 | 0.0193 |
|  |  | GO:0016301 | kinase activity | 324 | 8 | 3.48 | 0.0213 |
|  |  | GO:0015291 | secondary active transmembrane transporter activity | 56 | 3 | 0.6 | 0.0217 |
|  |  | GO:0000981 | DNA-binding transcription factor activity, RNA polymerase II-specific | 23 | 2 | 0.25 | 0.0248 |
|  |  | GO:0140612 | DNA damage sensor activity | 25 | 2 | 0.27 | 0.0290 |
|  |  | GO:0140664 | ATP-dependent DNA damage sensor activity | 25 | 2 | 0.27 | 0.0290 |
|  |  | GO:0140299 | small molecule sensor activity | 25 | 2 | 0.27 | 0.0290 |
|  |  | GO:0020037 | heme binding | 63 | 3 | 0.68 | 0.0296 |
|  |  | GO:0046906 | tetrapyrrole binding | 63 | 3 | 0.68 | 0.0296 |
|  |  | GO:0016874 | ligase activity | 68 | 3 | 0.73 | 0.0359 |
|  |  | GO:0016798 | hydrolase activity, acting on glycosyl bonds | 71 | 3 | 0.76 | 0.0401 |
|  |  | GO:0004553 | hydrolase activity, hydrolyzing O-glycosyl compounds | 71 | 3 | 0.76 | 0.0401 |
|  |  | GO:0004176 | ATP-dependent peptidase activity | 33 | 2 | 0.35 | 0.0483 |
|  | **CC** | GO:0030687 | preribosome, large subunit precursor | 15 | 2 | 0.08 | 0.0027 |
|  |  | GO:0005667 | transcription regulator complex | 15 | 2 | 0.08 | 0.0027 |
| **Nitrate** | **BP** | GO:0035269 | protein O-linked mannosylation | 14 | 3 | 0.18 | 0.0007 |
|  |  | GO:0042255 | ribosome assembly | 16 | 3 | 0.21 | 0.0010 |
|  |  | GO:0022618 | protein-RNA complex assembly | 18 | 3 | 0.24 | 0.0015 |
|  |  | GO:0010498 | proteasomal protein catabolic process | 71 | 5 | 0.93 | 0.0020 |
|  |  | GO:0006996 | organelle organization | 27 | 5 | 0.35 | 0.0074 |
|  |  | GO:0034976 | response to endoplasmic reticulum stress | 36 | 3 | 0.47 | 0.0110 |
|  |  | GO:0042273 | ribosomal large subunit biogenesis | 13 | 2 | 0.17 | 0.0119 |
|  |  | GO:0000027 | ribosomal large subunit assembly | 13 | 2 | 0.17 | 0.0119 |
|  |  | GO:0043161 | proteasome-mediated ubiquitin-dependent protein catabolic process | 41 | 3 | 0.54 | 0.0157 |
|  |  | GO:0010033 | response to organic substance | 43 | 3 | 0.56 | 0.0178 |
|  |  | GO:0006913 | nucleocytoplasmic transport | 17 | 2 | 0.22 | 0.0200 |
|  |  | GO:0051168 | nuclear export | 17 | 2 | 0.22 | 0.0200 |
|  |  | GO:0051169 | nuclear transport | 17 | 2 | 0.22 | 0.0200 |
|  |  | GO:0019752 | carboxylic acid metabolic process | 48 | 3 | 0.63 | 0.0239 |
|  |  | GO:0006082 | organic acid metabolic process | 48 | 3 | 0.63 | 0.0239 |
|  |  | GO:0043436 | oxoacid metabolic process | 48 | 3 | 0.63 | 0.0239 |
|  |  | GO:0006631 | fatty acid metabolic process | 20 | 2 | 0.26 | 0.0273 |
|  |  | GO:0006635 | fatty acid beta-oxidation | 20 | 2 | 0.26 | 0.0273 |
|  |  | GO:0016042 | lipid catabolic process | 20 | 2 | 0.26 | 0.0273 |
|  |  | GO:0044242 | cellular lipid catabolic process | 20 | 2 | 0.26 | 0.0273 |
|  |  | GO:0030258 | lipid modification | 20 | 2 | 0.26 | 0.0273 |
|  |  | GO:0034440 | lipid oxidation | 20 | 2 | 0.26 | 0.0273 |
|  |  | GO:0016054 | organic acid catabolic process | 20 | 2 | 0.26 | 0.0273 |
|  |  | GO:0009062 | fatty acid catabolic process | 20 | 2 | 0.26 | 0.0273 |
|  |  | GO:0044255 | cellular lipid metabolic process | 20 | 2 | 0.26 | 0.0273 |
|  |  | GO:0019395 | fatty acid oxidation | 20 | 2 | 0.26 | 0.0273 |
|  |  | GO:0046395 | carboxylic acid catabolic process | 20 | 2 | 0.26 | 0.0273 |
|  |  | GO:0072329 | monocarboxylic acid catabolic process | 20 | 2 | 0.26 | 0.0273 |
|  |  | GO:0044281 | small molecule metabolic process | 55 | 3 | 0.72 | 0.0340 |
|  |  | GO:0044282 | small molecule catabolic process | 23 | 2 | 0.3 | 0.0355 |
|  |  | GO:0032787 | monocarboxylic acid metabolic process | 23 | 2 | 0.3 | 0.0355 |
|  | **MF** | GO:0015020 | glucuronosyltransferase activity | 11 | 2 | 0.09 | 0.0033 |
|  |  | GO:0042285 | xylosyltransferase activity | 11 | 2 | 0.09 | 0.0033 |
|  |  | GO:0016747 | acyltransferase activity, transferring groups other than amino-acyl groups | 82 | 4 | 0.66 | 0.0040 |
|  |  | GO:0003824 | catalytic activity | 2264 | 30 | 18.21 | 0.0042 |
|  |  | GO:0016787 | hydrolase activity | 667 | 11 | 5.37 | 0.0144 |
|  |  | GO:0000049 | tRNA binding | 37 | 2 | 0.3 | 0.0353 |
|  |  | GO:0016887 | ATP hydrolysis activity | 233 | 5 | 1.87 | 0.0378 |
|  |  | GO:0017111 | ribonucleoside triphosphate phosphatase activity | 320 | 6 | 2.57 | 0.0414 |
|  |  | GO:0016817 | hydrolase activity, acting on acid anhydrides | 320 | 6 | 2.57 | 0.0414 |
|  |  | GO:0016818 | hydrolase activity, acting on acid anhydrides, in phosphorus-containing anhydrides | 320 | 6 | 2.57 | 0.0414 |
|  |  | GO:0016462 | pyrophosphatase activity | 320 | 6 | 2.57 | 0.0414 |
|  |  | GO:0016740 | transferase activity | 888 | 16 | 7.14 | 0.0461 |
|  |  | GO:0004674 | protein serine/threonine kinase activity | 171 | 4 | 1.38 | 0.0475 |
|  | **CC** | GO:0043231 | intracellular membrane-bounded organelle | 1501 | 20 | 12.64 | 0.0056 |
|  |  | GO:0030687 | preribosome, large subunit precursor | 15 | 2 | 0.13 | 0.0067 |
|  |  | GO:0005839 | proteasome core complex | 21 | 2 | 0.18 | 0.0130 |
|  |  | GO:0000502 | proteasome complex | 21 | 2 | 0.18 | 0.0130 |
|  |  | GO:1905368 | peptidase complex | 21 | 2 | 0.18 | 0.0130 |
|  |  | GO:1905369 | endopeptidase complex | 21 | 2 | 0.18 | 0.0130 |
|  |  | GO:1902494 | catalytic complex | 63 | 3 | 0.53 | 0.0152 |
|  |  | GO:0140535 | intracellular protein-containing complex | 25 | 2 | 0.21 | 0.0182 |
|  |  | GO:0031975 | envelope | 68 | 3 | 0.57 | 0.0187 |
|  |  | GO:0031967 | organelle envelope | 68 | 3 | 0.57 | 0.0187 |
|  |  | GO:0032991 | protein-containing complex | 225 | 7 | 1.89 | 0.0223 |

**Table S6 – MSMS data and IDs matched to m/z peaks of interest.** Confidence level assigned with *denoting m/z match or fragment match and ** denoting m/z and fragmentation match.

| ***Precursor ion m/z* (Da)** | ***Product ions* (Da)** | **Proposed ID** | ***m/z* error (Da)** | **Confidence** |
| --- | --- | --- | --- | --- |
| 173.0156 | 112.991,154.952, 89.024, 61.988, 78.959 | Phenyl dihydrogen phosphate | 0.02 | ** |
| 277.2154 |  | FA (18:3),  α-linolenic acid | 0.001 | * |
| 481.2407 |  | LPG 16:1 | 0.02 | * |
| 505.2492 | 277.222, 253.100, 168.031, 341.110, 441.141, 152.995 | LPG 18:3 | 0.007 | ** |
| 741.4446 | 505.264, 277.222, 253.223, 733.516, 153.000 | PG 34:4 | 0.01 | ** |
| 745.4753 | 255.238, 279.236, 507.283, 152.995, 415.232 | PG 34:2 | 0.02 | ** |
| 765.4570 | 255.238, 279.236, 152.995, 519.247, 505.264, 415.232, 693.461 | PG 36:6 | 0.02 | ** |
| 767.4688 |  | PG 36:5 | 0.02 | * |
| 773.4932 | 483.280, 255.238, 153.000, 537.249 | PG 36:2 | 0.04 | * |
| 791.4698 | 537.282, 225.009, 255.232,164.987 | SQDG 32:1 | 0.03 | ** |
| 793.4842 | 225.009, 537.282, 255.238 | SQDG 32:0 | 0.03 | ** |
| 817.4826 | 225.009, 555.293, 164.987, 537.274, 355.073 | SQDG 34:2 | 0.03 | ** |
| 833.4860 | 255.232, 279.236, 241.015, 553.287, 391.235,152.995, 571.301 | PI 34:2 | 0.03 | ** |
| 837.456 |  | SQDG 36:6 | 0.03 | * |
| 839.4663 | 225.009, 255.238, 153.030, 721.391, 761.427, 801.467 | SQDG 36:5 | 0.03 | ** |
| 841.4805 | 225.009, 279.236, 561.290,164.992, 255.238 | SQDG 36:4 | 0.03 | ** |
| 849.4737 | 721.391, 761.427, 241.015, 577.283, 295.236, 153.000 | PI 36:8 | 0.02 | ** |
| 855.4684 | 279.236, 241.015, 152.995, 577.283, 415.232 | PI 36:5 | 0.03 | * |
| 857.4848 | 279.236, 241.015, 577.291, 415.232,152.995 | PI 36:4 | 0.03 | ** |
| 907.5068 |  | PI 40:7 | 0.03 | * |
| 909.5186 |  | PI 40:6 | 0.03 | * |
| 1375.7160 | 745.502, 719.492, 255.232, 279.230, 627.208, 415.225, 483.272,152.991 | CL 66:2 | 0.2 | * |

**Fig S6 – Mean intensity over time for peaks of interest with ID in negative mode.**

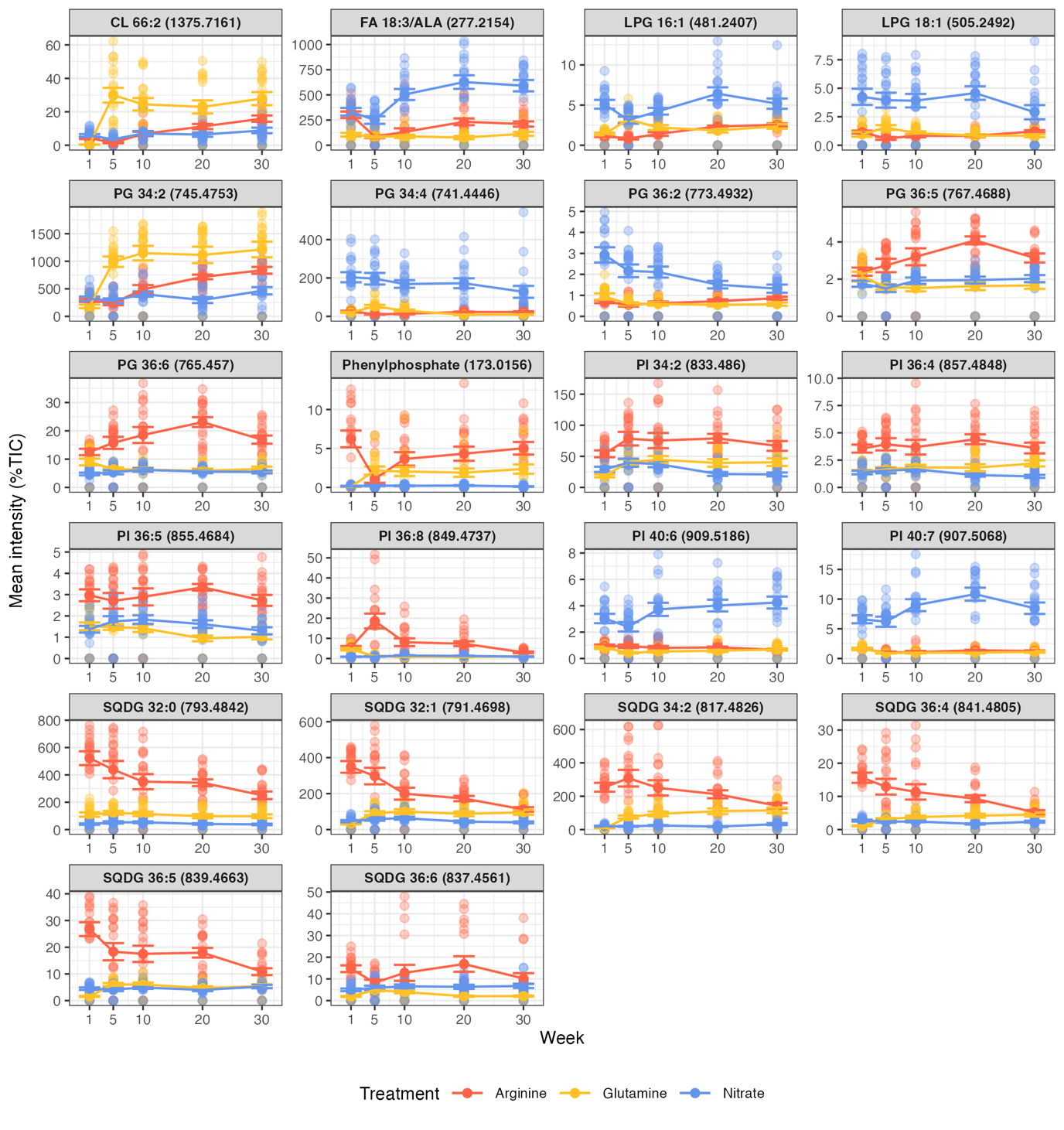

**Table S7 – Linear model output for peaks of interest with confirmed IDs.** Test statistics reported are primarily from mixed effects models fit with lmer() with biological replicate as the random effect. Where this resulted in a singular fit, a linear model without the random effect was fit instead.

| ***m/z*** | **effect** | **χ^2^/F** | **df** | **p** | **Model** |
| --- | --- | --- | --- | --- | --- |
| 173.0156 | Treatment | 511.28 | 2 | <0.001 | LMM |
|  | Week | 9.42 | 4 | 0.051 | LMM |
|  | Treatment:Week | 97.49 | 8 | <0.001 | LMM |
| 277.2154 | Treatment | 526.67 | 2 | <0.001 | LMM |
|  | Week | 49.14 | 4 | <0.001 | LMM |
|  | Treatment:Week | 67.28 | 8 | <0.001 | LMM |
| 481.2407 | Treatment | 845.01 | 2 | <0.001 | LMM |
|  | Week | 88.7 | 4 | <0.001 | LMM |
|  | Treatment:Week | 162.43 | 8 | <0.001 | LMM |
| 505.2492 | Treatment | 601.68 | 2 | <0.001 | LMM |
|  | Week | 36.5 | 4 | <0.001 | LMM |
|  | Treatment:Week | 64.3 | 8 | <0.001 | LMM |
| 741.4446 | Treatment | 899.29 | 2 | <0.001 | LMM |
|  | Week | 74.14 | 4 | <0.001 | LMM |
|  | Treatment:Week | 176.08 | 8 | <0.001 | LMM |
| 745.4753 | Treatment | 409.57 | 2 | <0.001 | LMM |
|  | Week | 389.78 | 4 | <0.001 | LMM |
|  | Treatment:Week | 262.18 | 8 | <0.001 | LMM |
| 765.4570 | Treatment | 728.53 | 2 | <0.001 | LMM |
|  | Week | 6.45 | 4 | 0.168 | LMM |
|  | Treatment:Week | 49.71 | 8 | <0.001 | LMM |
| 767.4688 | Treatment | 222.36 | 2, 217 | <0.001 | LM |
|  | Week | 4.11 | 4, 217 | 0.003 | LM |
|  | Treatment:Week | 11.83 | 8, 217 | <0.001 | LM |
| 773.4932 | Treatment | 313.62 | 2, 217 | <0.001 | LM |
|  | Week | 10.34 | 4, 217 | <0.001 | LM |
|  | Treatment:Week | 10.86 | 8, 217 | <0.001 | LM |
| 791.4698 | Treatment | 284.9 | 2, 217 | <0.001 | LM |
|  | Week | 16.93 | 4, 217 | <0.001 | LM |
|  | Treatment:Week | 19.76 | 8, 217 | <0.001 | LM |
| 793.4842 | Treatment | 742.01 | 2, 217 | <0.001 | LM |
|  | Week | 13.53 | 4, 217 | <0.001 | LM |
|  | Treatment:Week | 2.88 | 8, 217 | 0.005 | LM |
| 817.4826 | Treatment | 576.45 | 2, 217 | <0.001 | LM |
|  | Week | 21.52 | 4, 217 | <0.001 | LM |
|  | Treatment:Week | 41.22 | 8, 217 | <0.001 | LM |
| 833.4860 | Treatment | 273.04 | 2 | <0.001 | LMM |
|  | Week | 69.52 | 4 | <0.001 | LMM |
|  | Treatment:Week | 47.81 | 8 | <0.001 | LMM |
| 837.4561 | Treatment | 273.01 | 2 | <0.001 | LMM |
|  | Week | 21.93 | 4 | <0.001 | LMM |
|  | Treatment:Week | 51.83 | 8 | <0.001 | LMM |
| 839.4663 | Treatment | 626.31 | 2 | <0.001 | LMM |
|  | Week | 39.9 | 4 | <0.001 | LMM |
|  | Treatment:Week | 143.53 | 8 | <0.001 | LMM |
| 841.4805 | Treatment | 896.04 | 2 | <0.001 | LMM |
|  | Week | 29.24 | 4 | <0.001 | LMM |
|  | Treatment:Week | 378.42 | 8 | <0.001 | LMM |
| 849.4737 | Treatment | 500.18 | 2 | <0.001 | LMM |
|  | Week | 46.84 | 4 | <0.001 | LMM |
|  | Treatment:Week | 188.04 | 8 | <0.001 | LMM |
| 855.4684 | Treatment | 538.84 | 2 | <0.001 | LMM |
|  | Week | 65.09 | 4 | <0.001 | LMM |
|  | Treatment:Week | 33.92 | 8 | <0.001 | LMM |
| 857.4848 | Treatment | 248.58 | 2 | <0.001 | LMM |
|  | Week | 8.89 | 4 | 0.064 | LMM |
|  | Treatment:Week | 32.88 | 8 | <0.001 | LMM |
| 907.5068 | Treatment | 787.52 | 2 | <0.001 | LMM |
|  | Week | 70.62 | 4 | <0.001 | LMM |
|  | Treatment:Week | 89.52 | 8 | <0.001 | LMM |
| 909.5186 | Treatment | 692.34 | 2 | <0.001 | LMM |
|  | Week | 14.02 | 4 | 0.007 | LMM |
|  | Treatment:Week | 87.93 | 8 | <0.001 | LMM |
| 1375.7160 | Treatment | 201.97 | 2 | <0.001 | LMM |
|  | Week | 270.75 | 4 | <0.001 | LMM |
|  | Treatment:Week | 307 | 8 | <0.001 | LMM |

**Table S8 – Pairwise contrasts with emmeans for peaks of interest with confirmed IDs with fold change (fc) and Log2-fold change (log2fc).**

| ***m/z*** | **Week** | **Contrast** | **t-ratio** | **df** | **p** | **fc** | **Log2fc** |
| --- | --- | --- | --- | --- | --- | --- | --- |
| **173.0156** | **1** | A—G | 12.47 | 133.2 | <0.001 | 32.7 | 5.03 |
|  |  | A—N | 12.94 | 133.3 | <0.001 | 28.5 | 4.83 |
|  |  | G—N | -0.8 | 134.1 | 0.425 | 0.87 | -0.2 |
|  | **5** | A—G | -0.01 | 133.1 | 0.991 | 1.02 | 0.03 |
|  |  | A—N | 4.18 | 132.3 | <0.001 | 6.13 | 2.62 |
|  |  | G—N | 4.93 | 133.3 | <0.001 | 6.01 | 2.59 |
|  | **10** | A—G | 3.76 | 133.6 | <0.001 | 1.79 | 0.84 |
|  |  | A—N | 10.11 | 132.8 | <0.001 | 19 | 4.25 |
|  |  | G—N | 6.26 | 132.5 | <0.001 | 10.6 | 3.41 |
|  | **20** | A—G | 2.87 | 133.6 | 0.005 | 1.52 | 0.6 |
|  |  | A—N | 11.16 | 133.1 | <0.001 | 18.7 | 4.22 |
|  |  | G—N | 7.04 | 132.8 | <0.001 | 12.3 | 3.62 |
|  | **30** | A—G | 3.27 | 132.9 | 0.001 | 1.68 | 0.74 |
|  |  | A—N | 10.39 | 133.5 | <0.001 | 20.8 | 4.38 |
|  |  | G—N | 7.27 | 132.7 | <0.001 | 12.4 | 3.63 |
| **277.2154** | **1** | A—G | 7.12 | 212.7 | <0.001 | 2.71 | 1.44 |
|  |  | A—N | -0.51 | 212.3 | 0.607 | 0.911 | -0.14 |
|  |  | G—N | -7.63 | 212.4 | <0.001 | 0.336 | -1.57 |
|  | **5** | A—G | 0.56 | 212.2 | 0.577 | 1.13 | 0.17 |
|  |  | A—N | -6.73 | 212.3 | <0.001 | 0.345 | -1.53 |
|  |  | G—N | -7.51 | 212.5 | <0.001 | 0.306 | -1.71 |
|  | **10** | A—G | 1.86 | 212.1 | 0.064 | 1.6 | 0.68 |
|  |  | A—N | -9.17 | 212.5 | <0.001 | 0.3 | -1.74 |
|  |  | G—N | -11.26 | 212.4 | <0.001 | 0.188 | -2.41 |
|  | **20** | A—G | 5.24 | 212.2 | <0.001 | 2.36 | 1.24 |
|  |  | A—N | -8.45 | 212.2 | <0.001 | 0.33 | -1.6 |
|  |  | G—N | -13.02 | 212.5 | <0.001 | 0.14 | -2.84 |
|  | **30** | A—G | 3.51 | 212.4 | <0.001 | 1.64 | 0.71 |
|  |  | A—N | -8.03 | 212.4 | <0.001 | 0.336 | -1.57 |
|  |  | G—N | -11.24 | 212.4 | <0.001 | 0.205 | -2.29 |
| **481.2407** | **1** | A—G | -2.47 | 212.8 | 0.014 | 0.759 | -0.4 |
|  |  | A—N | -17.17 | 212.3 | <0.001 | 0.225 | -2.15 |
|  |  | G—N | -14.41 | 212.5 | <0.001 | 0.296 | -1.76 |
|  | **5** | A—G | -12.68 | 212.2 | <0.001 | 0.297 | -1.75 |
|  |  | A—N | -13.57 | 212.3 | <0.001 | 0.267 | -1.91 |
|  |  | G—N | -1.33 | 212.6 | 0.183 | 0.899 | -0.15 |
|  | **10** | A—G | -4.17 | 212.1 | <0.001 | 0.684 | -0.55 |
|  |  | A—N | -12.06 | 212.6 | <0.001 | 0.38 | -1.39 |
|  |  | G—N | -7.97 | 212.5 | <0.001 | 0.556 | -0.85 |
|  | **20** | A—G | 0 | 212.3 | 0.998 | 0.984 | -0.02 |
|  |  | A—N | -12.26 | 212.2 | <0.001 | 0.325 | -1.62 |
|  |  | G—N | -11.5 | 212.5 | <0.001 | 0.331 | -1.6 |
|  | **30** | A—G | -0.61 | 212.5 | 0.541 | 0.95 | -0.07 |
|  |  | A—N | -8.92 | 212.4 | <0.001 | 0.462 | -1.11 |
|  |  | G—N | -8.04 | 212.5 | <0.001 | 0.486 | -1.04 |
| **505.2492** | **1** | A—G | -2.38 | 186.7 | 0.018 | 0.803 | -0.32 |
|  |  | A—N | -8.89 | 185.2 | <0.001 | 0.236 | -2.08 |
|  |  | G—N | -5.69 | 185.7 | <0.001 | 0.294 | -1.77 |
|  | **5** | A—G | -7.13 | 185.5 | <0.001 | 0.525 | -0.93 |
|  |  | A—N | -11.99 | 186.2 | <0.001 | 0.18 | -2.47 |
|  |  | G—N | -5.51 | 185.3 | <0.001 | 0.343 | -1.54 |
|  | **10** | A—G | -3.12 | 184.8 | 0.002 | 0.776 | -0.37 |
|  |  | A—N | -11.54 | 185.4 | <0.001 | 0.212 | -2.24 |
|  |  | G—N | -8.43 | 184.8 | <0.001 | 0.273 | -1.87 |
|  | **20** | A—G | 0.71 | 186.4 | 0.478 | 1.06 | 0.08 |
|  |  | A—N | -11.13 | 185.9 | <0.001 | 0.211 | -2.25 |
|  |  | G—N | -12.1 | 185 | <0.001 | 0.2 | -2.32 |
|  | **30** | A—G | 2.06 | 184.6 | 0.041 | 1.19 | 0.25 |
|  |  | A—N | -8.44 | 185.5 | <0.001 | 0.297 | -1.75 |
|  |  | G—N | -9.98 | 185.5 | <0.001 | 0.249 | -2 |
| **741.4446** | **1** | A—G | 2.07 | 212.3 | 0.04 | 1.21 | 0.28 |
|  |  | A—N | -9.61 | 212.1 | <0.001 | 0.131 | -2.94 |
|  |  | G—N | -11.52 | 212.2 | <0.001 | 0.108 | -3.21 |
|  | **5** | A—G | -9.48 | 212.1 | <0.001 | 0.229 | -2.13 |
|  |  | A—N | -15.87 | 212.1 | <0.001 | 0.053 | -4.24 |
|  |  | G—N | -6.91 | 212.2 | <0.001 | 0.232 | -2.11 |
|  | **10** | A—G | -4.91 | 212.1 | <0.001 | 0.485 | -1.04 |
|  |  | A—N | -13.49 | 212.2 | <0.001 | 0.0911 | -3.46 |
|  |  | G—N | -8.67 | 212.2 | <0.001 | 0.188 | -2.41 |
|  | **20** | A—G | 5.25 | 212.1 | <0.001 | 2.11 | 1.07 |
|  |  | A—N | -11.6 | 212.1 | <0.001 | 0.118 | -3.08 |
|  |  | G—N | -15.98 | 212.2 | <0.001 | 0.0561 | -4.16 |
|  | **30** | A—G | 4.96 | 212.2 | <0.001 | 2.17 | 1.12 |
|  |  | A—N | -9.54 | 212.2 | <0.001 | 0.168 | -2.58 |
|  |  | G—N | -14.13 | 212.2 | <0.001 | 0.0773 | -3.69 |
| **745.4753** | **1** | A—G | 2.61 | 212.2 | 0.01 | 1.51 | 0.59 |
|  |  | A—N | -0.93 | 212.1 | 0.355 | 0.912 | -0.13 |
|  |  | G—N | -3.52 | 212.1 | <0.001 | 0.605 | -0.72 |
|  | **5** | A—G | -12.47 | 212.1 | <0.001 | 0.279 | -1.84 |
|  |  | A—N | -1.22 | 212.1 | 0.222 | 0.863 | -0.21 |
|  |  | G—N | 11.2 | 212.1 | <0.001 | 3.09 | 1.63 |
|  | **10** | A—G | -7.87 | 212 | <0.001 | 0.457 | -1.13 |
|  |  | A—N | 2.92 | 212.1 | 0.004 | 1.38 | 0.47 |
|  |  | G—N | 11.11 | 212.1 | <0.001 | 3.03 | 1.6 |
|  | **20** | A—G | -7.32 | 212.1 | <0.001 | 0.499 | -1 |
|  |  | A—N | 8.26 | 212.1 | <0.001 | 2.15 | 1.1 |
|  |  | G—N | 14.87 | 212.1 | <0.001 | 4.31 | 2.11 |
|  | **30** | A—G | -5.19 | 212.1 | <0.001 | 0.608 | -0.72 |
|  |  | A—N | 5.75 | 212.1 | <0.001 | 1.7 | 0.76 |
|  |  | G—N | 10.69 | 212.1 | <0.001 | 2.79 | 1.48 |
| **765.4570** | **1** | A—G | 2.99 | 213 | 0.003 | 1.3 | 0.38 |
|  |  | A—N | 12.12 | 212.4 | <0.001 | 2.64 | 1.4 |
|  |  | G—N | 8.93 | 212.6 | <0.001 | 2.02 | 1.01 |
|  | **5** | A—G | 9.48 | 212.3 | <0.001 | 2.78 | 1.47 |
|  |  | A—N | 10.44 | 212.5 | <0.001 | 2.92 | 1.55 |
|  |  | G—N | 1.3 | 212.7 | 0.193 | 1.05 | 0.07 |
|  | **10** | A—G | 9.71 | 212.2 | <0.001 | 3.17 | 1.67 |
|  |  | A—N | 11.09 | 212.7 | <0.001 | 3.41 | 1.77 |
|  |  | G—N | 1.26 | 212.6 | 0.208 | 1.08 | 0.11 |
|  | **20** | A—G | 9.53 | 212.4 | <0.001 | 3.07 | 1.62 |
|  |  | A—N | 13.27 | 212.3 | <0.001 | 3.67 | 1.88 |
|  |  | G—N | 3.18 | 212.7 | 0.002 | 1.2 | 0.26 |
|  | **30** | A—G | 7.65 | 212.7 | <0.001 | 2.27 | 1.19 |
|  |  | A—N | 12.14 | 212.6 | <0.001 | 2.9 | 1.54 |
|  |  | G—N | 4.22 | 212.6 | <0.001 | 1.28 | 0.35 |
| **767.4688** | **1** | A—G | 1.14 | 217 | 0.257 | 1.05 | 0.07 |
|  |  | A—N | 6.37 | 217 | <0.001 | 1.37 | 0.45 |
|  |  | G—N | 5.13 | 217 | <0.001 | 1.3 | 0.38 |
|  | **5** | A—G | 11.67 | 217 | <0.001 | 1.97 | 0.98 |
|  |  | A—N | 9.82 | 217 | <0.001 | 1.81 | 0.86 |
|  |  | G—N | -1.53 | 217 | 0.128 | 0.921 | -0.12 |
|  | **10** | A—G | 11.83 | 217 | <0.001 | 2.25 | 1.17 |
|  |  | A—N | 8.22 | 217 | <0.001 | 1.91 | 0.93 |
|  |  | G—N | -3.87 | 217 | <0.001 | 0.846 | -0.24 |
|  | **20** | A—G | 9.38 | 217 | <0.001 | 1.95 | 0.96 |
|  |  | A—N | 8.77 | 217 | <0.001 | 1.86 | 0.89 |
|  |  | G—N | -0.9 | 217 | 0.372 | 0.954 | -0.07 |
|  | **30** | A—G | 8.66 | 217 | <0.001 | 1.66 | 0.73 |
|  |  | A—N | 5.75 | 217 | <0.001 | 1.46 | 0.54 |
|  |  | G—N | -2.96 | 217 | 0.003 | 0.875 | -0.19 |
| **773.4932** | **1** | A—G | -6.47 | 203 | <0.001 | 0.596 | -0.75 |
|  |  | A—N | -12.84 | 203 | <0.001 | 0.223 | -2.16 |
|  |  | G—N | -5.8 | 203 | <0.001 | 0.374 | -1.42 |
|  | **5** | A—G | 0.63 | 203 | 0.531 | 1.07 | 0.1 |
|  |  | A—N | -11.89 | 203 | <0.001 | 0.278 | -1.85 |
|  |  | G—N | -12.96 | 203 | <0.001 | 0.259 | -1.95 |
|  | **10** | A—G | 1.03 | 203 | 0.305 | 1.05 | 0.07 |
|  |  | A—N | -9.62 | 203 | <0.001 | 0.32 | -1.65 |
|  |  | G—N | -10.67 | 203 | <0.001 | 0.304 | -1.72 |
|  | **20** | A—G | 3.45 | 203 | <0.001 | 1.22 | 0.29 |
|  |  | A—N | -7.22 | 203 | <0.001 | 0.48 | -1.06 |
|  |  | G—N | -10.54 | 203 | <0.001 | 0.392 | -1.35 |
|  | **30** | A—G | 3.6 | 203 | <0.001 | 1.23 | 0.3 |
|  |  | A—N | -5.96 | 203 | <0.001 | 0.542 | -0.88 |
|  |  | G—N | -9.14 | 203 | <0.001 | 0.44 | -1.19 |
| **791.4698** | **1** | A—G | 16.09 | 217 | <0.001 | 9.78 | 3.29 |
|  |  | A—N | 14.51 | 217 | <0.001 | 7.46 | 2.9 |
|  |  | G—N | -1.82 | 217 | 0.071 | 0.763 | -0.39 |
|  | **5** | A—G | 7.65 | 217 | <0.001 | 3.71 | 1.89 |
|  |  | A—N | 10.21 | 217 | <0.001 | 5.07 | 2.34 |
|  |  | G—N | 2.9 | 217 | 0.004 | 1.37 | 0.45 |
|  | **10** | A—G | 4.41 | 217 | <0.001 | 2.14 | 1.1 |
|  |  | A—N | 9.26 | 217 | <0.001 | 3.62 | 1.86 |
|  |  | G—N | 4.88 | 217 | <0.001 | 1.69 | 0.76 |
|  | **20** | A—G | 2.95 | 217 | 0.003 | 1.53 | 0.62 |
|  |  | A—N | 10.91 | 217 | <0.001 | 3.49 | 1.8 |
|  |  | G—N | 7.36 | 217 | <0.001 | 2.28 | 1.19 |
|  | **30** | A—G | -0.01 | 217 | 0.993 | 1.03 | 0.05 |
|  |  | A—N | 8.31 | 217 | <0.001 | 2.64 | 1.4 |
|  |  | G—N | 8.06 | 217 | <0.001 | 2.55 | 1.35 |
| **793.4842** | **1** | A—G | 11.41 | 217 | <0.001 | 4.45 | 2.15 |
|  |  | A—N | 20.9 | 217 | <0.001 | 13 | 3.7 |
|  |  | G—N | 9.16 | 217 | <0.001 | 2.92 | 1.55 |
|  | **5** | A—G | 10.02 | 217 | <0.001 | 4.08 | 2.03 |
|  |  | A—N | 16.11 | 217 | <0.001 | 8.37 | 3.07 |
|  |  | G—N | 6.62 | 217 | <0.001 | 2.05 | 1.04 |
|  | **10** | A—G | 8.18 | 217 | <0.001 | 3.27 | 1.71 |
|  |  | A—N | 15.78 | 217 | <0.001 | 7.5 | 2.91 |
|  |  | G—N | 7.61 | 217 | <0.001 | 2.29 | 1.2 |
|  | **20** | A—G | 7.8 | 217 | <0.001 | 2.72 | 1.44 |
|  |  | A—N | 17.25 | 217 | <0.001 | 7.42 | 2.89 |
|  |  | G—N | 8.6 | 217 | <0.001 | 2.73 | 1.45 |
|  | **30** | A—G | 6.93 | 217 | <0.001 | 2.26 | 1.18 |
|  |  | A—N | 16.05 | 217 | <0.001 | 6.26 | 2.65 |
|  |  | G—N | 8.72 | 217 | <0.001 | 2.77 | 1.47 |
| **817.4826** | **1** | A—G | 21.85 | 217 | <0.001 | 27 | 4.75 |
|  |  | A—N | 17.6 | 217 | <0.001 | 12.3 | 3.63 |
|  |  | G—N | -4.53 | 217 | <0.001 | 0.458 | -1.13 |
|  | **5** | A—G | 11.24 | 217 | <0.001 | 4.68 | 2.23 |
|  |  | A—N | 17.29 | 217 | <0.001 | 16 | 4 |
|  |  | G—N | 6.61 | 217 | <0.001 | 3.42 | 1.78 |
|  | **10** | A—G | 7.52 | 217 | <0.001 | 2.84 | 1.5 |
|  |  | A—N | 15.42 | 217 | <0.001 | 11.6 | 3.54 |
|  |  | G—N | 7.93 | 217 | <0.001 | 4.1 | 2.04 |
|  | **20** | A—G | 2.81 | 217 | 0.005 | 1.49 | 0.58 |
|  |  | A—N | 16.13 | 217 | <0.001 | 10.6 | 3.4 |
|  |  | G—N | 12.41 | 217 | <0.001 | 7.07 | 2.82 |
|  | **30** | A—G | 0.56 | 217 | 0.577 | 1.09 | 0.13 |
|  |  | A—N | 9.61 | 217 | <0.001 | 4.1 | 2.04 |
|  |  | G—N | 8.76 | 217 | <0.001 | 3.76 | 1.91 |
| **833.4860** | **1** | A—G | 7.97 | 213 | <0.001 | 2.73 | 1.45 |
|  |  | A—N | 5.71 | 212.4 | <0.001 | 1.86 | 0.9 |
|  |  | G—N | -2.36 | 212.6 | 0.019 | 0.682 | -0.55 |
|  | **5** | A—G | 6.53 | 212.3 | <0.001 | 2.21 | 1.14 |
|  |  | A—N | 5.67 | 212.4 | <0.001 | 1.97 | 0.98 |
|  |  | G—N | -0.67 | 212.7 | 0.503 | 0.895 | -0.16 |
|  | **10** | A—G | 4.38 | 212.2 | <0.001 | 1.81 | 0.86 |
|  |  | A—N | 6.68 | 212.7 | <0.001 | 2.29 | 1.2 |
|  |  | G—N | 2.28 | 212.6 | 0.024 | 1.26 | 0.34 |
|  | **20** | A—G | 3.6 | 212.3 | <0.001 | 1.56 | 0.64 |
|  |  | A—N | 9.32 | 212.3 | <0.001 | 3.25 | 1.7 |
|  |  | G—N | 5.24 | 212.7 | <0.001 | 2.08 | 1.06 |
|  | **30** | A—G | 2.98 | 212.7 | 0.003 | 1.45 | 0.54 |
|  |  | A—N | 8.3 | 212.6 | <0.001 | 3 | 1.59 |
|  |  | G—N | 5.1 | 212.6 | <0.001 | 2.07 | 1.05 |
| **837.4561** | **1** | A—G | 12.11 | 212.3 | <0.001 | 7.06 | 2.82 |
|  |  | A—N | 7.06 | 212.1 | <0.001 | 2.95 | 1.56 |
|  |  | G—N | -5.17 | 212.2 | <0.001 | 0.418 | -1.26 |
|  | **5** | A—G | 4.35 | 212.1 | <0.001 | 1.92 | 0.94 |
|  |  | A—N | 2.02 | 212.1 | 0.045 | 1.4 | 0.49 |
|  |  | G—N | -2.26 | 212.2 | 0.025 | 0.729 | -0.46 |
|  | **10** | A—G | 4.72 | 212 | <0.001 | 3.44 | 1.78 |
|  |  | A—N | 1.8 | 212.2 | 0.074 | 2.22 | 1.15 |
|  |  | G—N | -3.05 | 212.2 | 0.003 | 0.645 | -0.63 |
|  | **20** | A—G | 8.36 | 212.1 | <0.001 | 6.4 | 2.68 |
|  |  | A—N | 2.45 | 212.1 | 0.015 | 2.35 | 1.23 |
|  |  | G—N | -5.83 | 212.2 | <0.001 | 0.367 | -1.45 |
|  | **30** | A—G | 7.07 | 212.2 | <0.001 | 4.3 | 2.1 |
|  |  | A—N | 0.32 | 212.1 | 0.75 | 1.42 | 0.51 |
|  |  | G—N | -6.66 | 212.2 | <0.001 | 0.331 | -1.59 |
| **839.4663** | **1** | A—G | 18.03 | 212.4 | <0.001 | 15.6 | 3.96 |
|  |  | A—N | 13.24 | 212.2 | <0.001 | 5.96 | 2.58 |
|  |  | G—N | -5.03 | 212.2 | <0.001 | 0.382 | -1.39 |
|  | **5** | A—G | 6.83 | 212.1 | <0.001 | 3.47 | 1.79 |
|  |  | A—N | 10.72 | 212.2 | <0.001 | 4.44 | 2.15 |
|  |  | G—N | 4.24 | 212.3 | <0.001 | 1.28 | 0.36 |
|  | **10** | A—G | 5.57 | 212.1 | <0.001 | 3.1 | 1.63 |
|  |  | A—N | 10.39 | 212.3 | <0.001 | 4.09 | 2.03 |
|  |  | G—N | 4.83 | 212.2 | <0.001 | 1.32 | 0.4 |
|  | **20** | A—G | 7.68 | 212.1 | <0.001 | 2.96 | 1.56 |
|  |  | A—N | 12.12 | 212.1 | <0.001 | 3.97 | 1.99 |
|  |  | G—N | 3.9 | 212.3 | <0.001 | 1.34 | 0.43 |
|  | **30** | A—G | 4.07 | 212.2 | <0.001 | 1.76 | 0.82 |
|  |  | A—N | 6.4 | 212.2 | <0.001 | 1.94 | 0.95 |
|  |  | G—N | 2.2 | 212.2 | 0.029 | 1.1 | 0.14 |
| **841.4805** | **1** | A—G | 21.54 | 212.5 | <0.001 | 13.1 | 3.71 |
|  |  | A—N | 14.47 | 212.2 | <0.001 | 5.74 | 2.52 |
|  |  | G—N | -7.32 | 212.3 | <0.001 | 0.439 | -1.19 |
|  | **5** | A—G | 11.2 | 212.1 | <0.001 | 4.7 | 2.23 |
|  |  | A—N | 12.78 | 212.2 | <0.001 | 5.42 | 2.44 |
|  |  | G—N | 2.01 | 212.3 | 0.046 | 1.15 | 0.21 |
|  | **10** | A—G | 7.06 | 212.1 | <0.001 | 3.24 | 1.69 |
|  |  | A—N | 14.02 | 212.3 | <0.001 | 5.33 | 2.41 |
|  |  | G—N | 6.99 | 212.3 | <0.001 | 1.65 | 0.72 |
|  | **20** | A—G | 3.25 | 212.1 | 0.001 | 1.72 | 0.78 |
|  |  | A—N | 16.59 | 212.1 | <0.001 | 4.83 | 2.27 |
|  |  | G—N | 12.4 | 212.3 | <0.001 | 2.8 | 1.49 |
|  | **30** | A—G | -1.01 | 212.3 | 0.312 | 1.02 | 0.03 |
|  |  | A—N | 8.54 | 212.2 | <0.001 | 2.07 | 1.05 |
|  |  | G—N | 9.27 | 212.3 | <0.001 | 2.02 | 1.02 |
| **849.4737** | **1** | A—G | 0.6 | 212.3 | 0.548 | 1.12 | 0.17 |
|  |  | A—N | 12.25 | 212.1 | <0.001 | 6.33 | 2.66 |
|  |  | G—N | 11.44 | 212.2 | <0.001 | 5.64 | 2.49 |
|  | **5** | A—G | 12.68 | 212.1 | <0.001 | 19.2 | 4.26 |
|  |  | A—N | 11.15 | 212.1 | <0.001 | 19 | 4.25 |
|  |  | G—N | -1.17 | 212.2 | 0.245 | 0.994 | -0.01 |
|  | **10** | A—G | 10.61 | 212.1 | <0.001 | 9.68 | 3.28 |
|  |  | A—N | 6.52 | 212.2 | <0.001 | 6.34 | 2.66 |
|  |  | G—N | -4.32 | 212.2 | <0.001 | 0.655 | -0.61 |
|  | **20** | A—G | 10.77 | 212.1 | <0.001 | 7.03 | 2.81 |
|  |  | A—N | 7.92 | 212.1 | <0.001 | 5.12 | 2.36 |
|  |  | G—N | -3.05 | 212.2 | 0.003 | 0.729 | -0.46 |
|  | **30** | A—G | 5.8 | 212.2 | <0.001 | 2.62 | 1.39 |
|  |  | A—N | 7.91 | 212.2 | <0.001 | 3.06 | 1.61 |
|  |  | G—N | 1.94 | 212.2 | 0.054 | 1.17 | 0.22 |
| **855.4684** | **1** | A—G | 7.7 | 212.3 | <0.001 | 1.84 | 0.88 |
|  |  | A—N | 9.96 | 212.1 | <0.001 | 2.15 | 1.11 |
|  |  | G—N | 2.09 | 212.2 | 0.038 | 1.17 | 0.23 |
|  | **5** | A—G | 8.85 | 212.1 | <0.001 | 2.09 | 1.06 |
|  |  | A—N | 5 | 212.1 | <0.001 | 1.55 | 0.64 |
|  |  | G—N | -3.68 | 212.2 | <0.001 | 0.744 | -0.43 |
|  | **10** | A—G | 9.16 | 212.1 | <0.001 | 2.18 | 1.12 |
|  |  | A—N | 7.25 | 212.2 | <0.001 | 1.8 | 0.85 |
|  |  | G—N | -2.08 | 212.2 | 0.038 | 0.828 | -0.27 |
|  | **20** | A—G | 12.72 | 212.1 | <0.001 | 2.75 | 1.46 |
|  |  | A—N | 8 | 212.1 | <0.001 | 1.84 | 0.88 |
|  |  | G—N | -4.87 | 212.2 | <0.001 | 0.669 | -0.58 |
|  | **30** | A—G | 10.35 | 212.2 | <0.001 | 2.36 | 1.24 |
|  |  | A—N | 8.53 | 212.2 | <0.001 | 1.97 | 0.98 |
|  |  | G—N | -1.94 | 212.2 | 0.054 | 0.835 | -0.26 |
| **857.4848** | **1** | A—G | 6.34 | 212.8 | <0.001 | 2.38 | 1.25 |
|  |  | A—N | 7.06 | 212.3 | <0.001 | 2.6 | 1.38 |
|  |  | G—N | 0.6 | 212.5 | 0.547 | 1.09 | 0.12 |
|  | **5** | A—G | 6.7 | 212.2 | <0.001 | 2.72 | 1.45 |
|  |  | A—N | 6.09 | 212.4 | <0.001 | 2.59 | 1.37 |
|  |  | G—N | -0.41 | 212.6 | 0.681 | 0.951 | -0.07 |
|  | **10** | A—G | 4.27 | 212.1 | <0.001 | 2.16 | 1.11 |
|  |  | A—N | 5.59 | 212.6 | <0.001 | 2.54 | 1.34 |
|  |  | G—N | 1.28 | 212.5 | 0.202 | 1.18 | 0.23 |
|  | **20** | A—G | 4.17 | 212.3 | <0.001 | 1.9 | 0.92 |
|  |  | A—N | 8.27 | 212.2 | <0.001 | 3.46 | 1.79 |
|  |  | G—N | 3.71 | 212.6 | <0.001 | 1.83 | 0.87 |
|  | **30** | A—G | 1.3 | 212.5 | 0.195 | 1.45 | 0.53 |
|  |  | A—N | 7.56 | 212.4 | <0.001 | 3.35 | 1.74 |
|  |  | G—N | 6.05 | 212.5 | <0.001 | 2.31 | 1.21 |
| 907.5068 | 1 | A—G | -0.71 | 212.9 | 0.479 | 0.923 | -0.11 |
|  |  | A—N | -4.89 | 212.4 | <0.001 | 0.243 | -2.04 |
|  |  | G—N | -4.1 | 212.5 | <0.001 | 0.263 | -1.93 |
|  | 5 | A—G | 4.35 | 212.3 | <0.001 | 1.28 | 0.36 |
|  |  | A—N | -9.26 | 212.4 | <0.001 | 0.166 | -2.59 |
|  |  | G—N | -13.91 | 212.6 | <0.001 | 0.13 | -2.95 |
|  | 10 | A—G | 0.89 | 212.2 | 0.376 | 1.18 | 0.24 |
|  |  | A—N | -12.71 | 212.6 | <0.001 | 0.136 | -2.88 |
|  |  | G—N | -13.86 | 212.5 | <0.001 | 0.116 | -3.11 |
|  | 20 | A—G | 1.25 | 212.3 | 0.212 | 1.09 | 0.12 |
|  |  | A—N | -13.27 | 212.3 | <0.001 | 0.113 | -3.14 |
|  |  | G—N | -13.67 | 212.6 | <0.001 | 0.104 | -3.26 |
|  | 30 | A—G | -0.05 | 212.6 | 0.961 | 0.984 | -0.02 |
|  |  | A—N | -11.4 | 212.5 | <0.001 | 0.14 | -2.84 |
|  |  | G—N | -11 | 212.5 | <0.001 | 0.142 | -2.81 |
| 909.5186 | 1 | A—G | 0.73 | 213.6 | 0.467 | 1.09 | 0.13 |
|  |  | A—N | -6.29 | 212.7 | <0.001 | 0.316 | -1.66 |
|  |  | G—N | -6.91 | 213 | <0.001 | 0.289 | -1.79 |
|  | 5 | A—G | 8.51 | 212.5 | <0.001 | 2.46 | 1.3 |
|  |  | A—N | -4.12 | 212.8 | <0.001 | 0.387 | -1.37 |
|  |  | G—N | -12.76 | 213.2 | <0.001 | 0.158 | -2.67 |
|  | 10 | A—G | 3.76 | 212.3 | <0.001 | 1.62 | 0.7 |
|  |  | A—N | -8.6 | 213.2 | <0.001 | 0.248 | -2.01 |
|  |  | G—N | -12.64 | 213 | <0.001 | 0.153 | -2.71 |
|  | 20 | A—G | 1.24 | 212.6 | 0.218 | 1.11 | 0.15 |
|  |  | A—N | -11.19 | 212.5 | <0.001 | 0.188 | -2.41 |
|  |  | G—N | -11.7 | 213.2 | <0.001 | 0.169 | -2.56 |
|  | 30 | A—G | -1.58 | 213.1 | 0.117 | 0.854 | -0.23 |
|  |  | A—N | -13.27 | 212.9 | <0.001 | 0.147 | -2.76 |
|  |  | G—N | -11.31 | 213 | <0.001 | 0.172 | -2.54 |
| 1375.7160 | 1 | A—G | 5.3 | 212.2 | <0.001 | 9.21 | 3.2 |
|  |  | A—N | -1.97 | 212.1 | 0.05 | 0.726 | -0.46 |
|  |  | G—N | -7.24 | 212.1 | <0.001 | 0.0788 | -3.66 |
|  | 5 | A—G | -14.53 | 212.1 | <0.001 | 0.0585 | -4.09 |
|  |  | A—N | -3.14 | 212.1 | 0.002 | 0.426 | -1.23 |
|  |  | G—N | 11.28 | 212.1 | <0.001 | 7.27 | 2.86 |
|  | 10 | A—G | -7.06 | 212 | <0.001 | 0.297 | -1.75 |
|  |  | A—N | 0.96 | 212.1 | 0.339 | 1.1 | 0.14 |
|  |  | G—N | 8.27 | 212.1 | <0.001 | 3.71 | 1.89 |
|  | 20 | A—G | -6.26 | 212.1 | <0.001 | 0.38 | -1.4 |
|  |  | A—N | 2.81 | 212.1 | 0.005 | 1.56 | 0.64 |
|  |  | G—N | 8.73 | 212.1 | <0.001 | 4.1 | 2.04 |
|  | 30 | A—G | -3.93 | 212.1 | <0.001 | 0.504 | -0.99 |
|  |  | A—N | 3.83 | 212.1 | <0.001 | 1.69 | 0.76 |
|  |  | G—N | 7.59 | 212.1 | <0.001 | 3.35 | 1.74 |

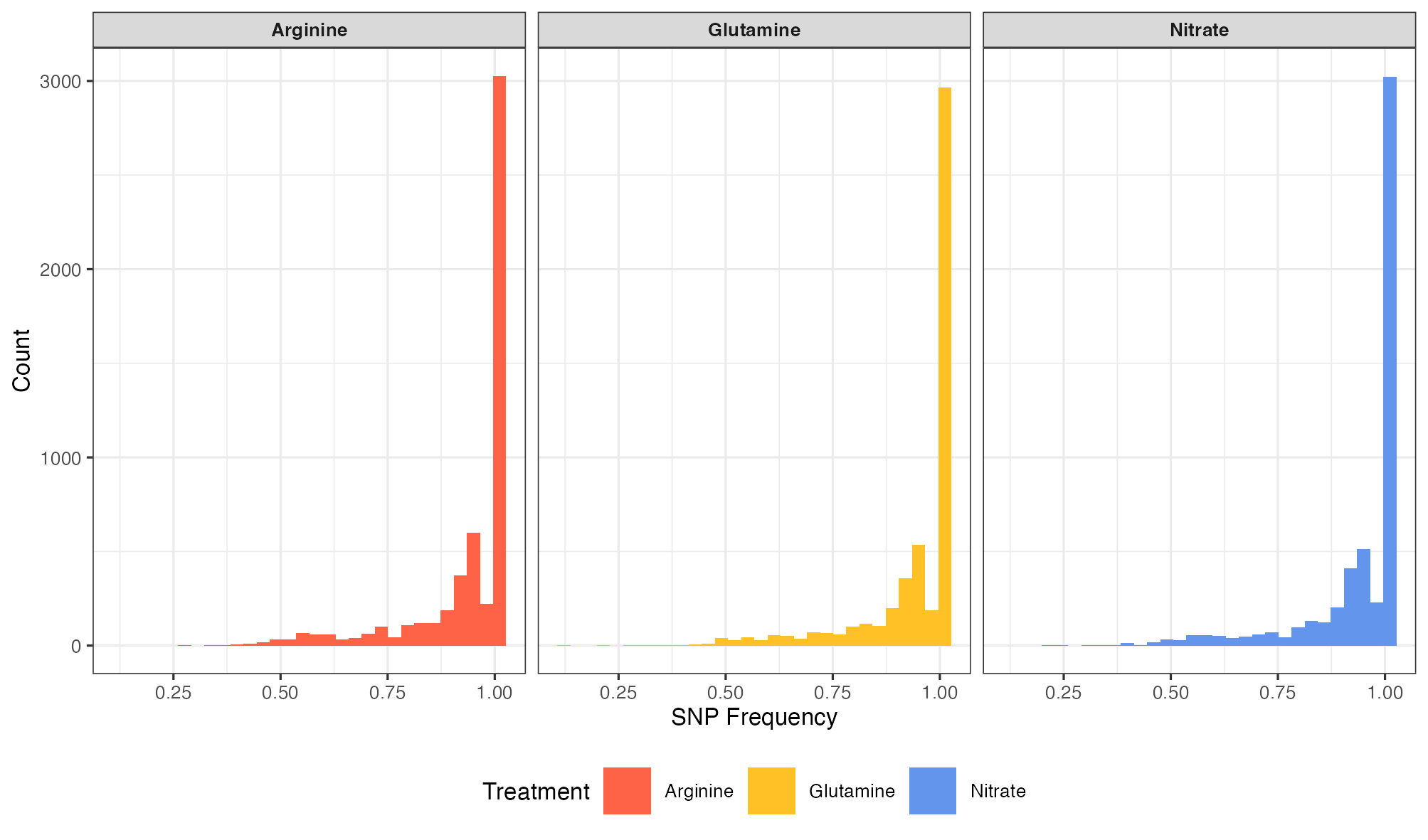

**Figure S7 – Distribution of SNP frequencies across treatments.**
